# Quantitative trait locus mapping of root exudate metabolome in a *Solanum lycopersicum* Moneymaker x *S. pimpinellifolium* RIL population and their putative links to rhizosphere microbiome

**DOI:** 10.64898/2025.12.17.693946

**Authors:** Bora Kim, Gertjan Kramer, Márcio Fernandes Alves Leite, Basten L. Snoek, Anouk Zancarini, Harro J. Bouwmeester

**Affiliations:** Plant Hormone Biology, Swammerdam Institute for Life Sciences, University of Amsterdam, 1098 XH Amsterdam, The Netherlands; Mass Spectrometry of Biomolecules, Swammerdam Institute for Life Sciences, University of Amsterdam, 1098 XH Amsterdam, The Netherlands; Theoretical Biology and Bioinformatics, Science for Life, Utrecht University, Padualaan 8, 3584 CH, Utrecht, The Netherlands; IGEPP, INRAE, Institut Agro, Univ Rennes, 35653, Le Rheu, France

**Keywords:** Root exudate, tomato, QTL, RIL, metabolomics

## Abstract

Crop resilience to abiotic and biotic stresses is vital for sustainable agriculture and global food security. The root exudates (RE)—a complex mixture of metabolites exuded by the plant roots—plays a central role in this resilience by mediating interactions with organisms around the roots, including soil microorganisms. These interactions influence nutrient cycling, the recruitment of beneficial microbes, and ultimately plant health. Although host genotype is known to affect the RE composition, the specific metabolites and underlying genetic mechanisms remain poorly understood. Here, we employed untargeted metabolomics to analyse the tomato RE composition and mapped individual metabolic features to the tomato genome using quantitative trait locus (QTL) analysis in a recombinant inbred line population derived from *Solanum lycopersicum* and *Solanum pimpinellifolium*. Our results shed light on the intricate chemical composition of the tomato RE, and reveal domestication-driven shifts in the RE composition and genetic loci associated with the abundance of specific metabolites. Notably, we identify overlap between metabolic and earlier published microbial QTLs, supportive of a link between RE composition and microbiome assembly, and providing a basis for further unravelling of the plant–microbe chemical interaction.

## Introduction

Crops are exposed to ceaseless abiotic and biotic stresses, which can greatly affect the quality and quantity of crop production. Therefore, crop resilience to such stresses is one of the most important challenges in agriculture, as it tightly links to global food security. Chemical fertilizers and pesticides are ubiquitously used to obtain satisfactory yield, however, increased public awareness of environmental protection results in tightening regulations on the use of chemicals in agriculture. A promising alternative to achieve a more sustainable agriculture, is to exploit the stress resilience related functions of plants that already exist in nature, one of which is the relationship with soil microorganisms.

The relationship between plants and soil microorganisms is particularly important around the roots, and plants steer this relationship through the exudation of small molecules from their roots into the surrounding rhizosphere soil. This collection of metabolites is called the root exudate (RE). The RE provides a carbon source for myriad of soil microbes, but also shapes the microbial community through a range of other mechanisms (Bouwmeester et al. 2025). Some of the microorganisms that gather around the roots can have a profound effect on plant health. They can promote plant growth by producing phytohormones and making soil nutrients available for plants. Moreover, microbes can increase plant immunity against pathogens and pests, or directly suppress pathogens by competing for niche space and nutrients, and by producing antimicrobial compounds (Vives-Peris et al. 2019). Interestingly, these, so-called beneficial microorganisms, are recruited through signaling molecules or are selected through antimicrobial compounds present in the RE. For instance, the signaling molecule strigolactones are actively secreted from roots under phosphate deficiency and attract arbuscular mycorrhizal fungi that help the plants absorb phosphate from the soil (Aliche et al. 2020). Due to the great impact of the root-associated microbiome on plant growth and health, there has been a strong increase in research to understand the mechanism underlying microbiome selection by plants.

Indeed, several authors shed light on the genetic basis underlying the recruitment of the root-associated microbiome using genetic association analyses. Genome Wide Association Studies identified host loci that correlate with the abundance of specific subsets of root-associated bacteria and fungi in different plant species (Bergelson *et al*., 2019; Deng *et al*., 2021; Sutherland *et al*., 2022). Moreover, quantitative trait locus (QTL) analysis with metagenomics data was done, and this revealed host genetic determinants of the microbiome and its functionality, and the effect of domestication on this process, in barley and tomato (Escudero-Martinez et al. 2022; Oyserman et al. 2022). It is highly likely that these host genetic determinants include the presence or abundance of specific metabolites in the root exudate (Bouwmeester et al. 2025), however, it remains an open question which compounds are exuded and how plant genetic factors regulate their release.

To answer these questions, insight is required into the metabolites that constitute the RE. Many studies have used targeted approaches to quantify the concentration of specific compounds in RE. These include organic acids, amino acids, sugars, fatty acids, vitamins and phenolics (Dennis *et al*., 2010). While this approach provides information about the RE, the targeted nature of the analysis, by definition, overlooks other, potentially unknown, compounds. A way around this pitfall is the use of metabolomics, an untargeted, sensitive and comprehensive analysis, of as many as possible small molecules. Over the past few decades, metabolomics has not only allowed to explore the diversity of plant metabolites, several studies already showed a great potential of metabolomics as a tool to investigate the relationship between root exudate and phenotypes, such as activity of root-knot nematode and Fe acquisition strategies in tomato and root-associated bacteria in *Avena barbata* (Yang *et al*., 2016; Zhalnina *et al*., 2018; Astolfi *et al*., 2020). In order to also find the genetic factors that drive possible changes in the RE composition, genetic association analyses can be employed. So far, only a few studies demonstrated the use of genetic studies to understand how specific rhizosphere signaling compounds - e.g. strigolactones, isoflavonoids, that are involved in the plant-microbe interaction - are related to the genetics of the plant host (Cardoso et al. 2014; Ramongolalaina, Teraishi and Okumoto 2018).

Therefore, in this study, we investigated the composition of tomato RE using metabolomics and explored the genetic basis of this composition in a RIL population derived from a cross between tomato (*Solanum lycopersicum* cv. Moneymaker) and its wild relative (*Solanum pimpinellifolium*) through QTL analysis. Metabolomics enabled us to capture the chemical diversity in the tomato RE and revealed the impact of domestication on its composition. Mapping subsequently allowed us to identify QTLs for RE metabolic features. Notably, some of these QTLs overlapped with QTLs for metagenomic contigs reported by Oyserman et al. (2022), suggesting a link between plant genetics, metabolite production, and rhizosphere microbial community composition.

## Materials and methods

### Root exudate collection

Seeds of a tomato recombinant inbred line (RIL) population, consisting of 100 genotypes derived from a cross between *S. lycopersicum* cv. Moneymaker (MM; LA2706) and *S. pimpinellifolium* (SP; CGN 15528, also named G1.1554), were provided by the Wageningen Seed Lab, Wageningen University, The Netherlands. The seeds (not surface-sterilized) were germinated on a greenhouse compost (Zaaigrond nr1+SIR, Jongkind, The Netherlands) at 22-25°C in 15/9 h day/night in the greenhouse facility, University of Amsterdam, The Netherlands. Then, the RIL seedlings were transplanted to 500 mL volume pot filled with 200 g of the compost (n = 5) and grown in the same environmental condition described above. The plant pots and pots without a plant (experimental blanks, n=5) were placed according to a random block design to avoid a local site effect on the plant growth. The pots with and without a plant were regularly watered with demineralized water to keep 80% water-holding capacity of the soil until root exudate (RE) collection. Twenty-eight days after transplanting, the pots were flushed with 5% ethanol until the extract volume reached 250 mL. Subsequently, the extract was centrifuged for 5 min at 4,300 rpm and the supernatant, which we coined RE, was stored at −20°C until further analysis.

### LC-ESI-QTOF-MS analysis

For the sample preparation for untargeted LC-MS analysis, pH of 25 mL RE sample was adjusted to 5 with 0.1 M HCl (in 5% ethanol) and loaded on solid phase extraction (SPE) cartridge (Discovery® DSC-18 500 mg/ 6 mL; Sigma-Aldrich, United States) which was pre-conditioned with 6 mL of 100 % acetone and 12 mL of 5% ethanol. Subsequently, the SPE cartridge was washed with 5% ethanol adjusted to pH 5. Then, the sample was eluted from the SPE cartridge with 3 mL acetone and dried in a vacuum concentrator (Scan Speed 40; Labogene, Lynge, Denmark) at 2,000 rpm for 2 h. Finally, the dried residue was reconstituted in 150 µL of 25 % acetonitrile and filtered through micro-centrifugal nylon filter (0.2 µm; Thermo Fisher Scientific, United States) in centrifuge for 1 min at 12,000 rpm. Aliquots of each sample were pooled as a quality control (QC) sample.

RE samples were analyzed using a QTOF MS equipped with a dual-stage trapped ion mobility separation cell (timsTOF pro Bruker Daltonics Inc, Billerica, MA). Sample injection (10 μL) and LC separation were performed on an Ultimate RS UHPLC system (Thermo scientific, Germeringen, Germany) with an Acquity UPLC CSH C18 130Å, 1.7 um, 2.1 mm × 100 mm protected by a VanGuard 2.1 mm x 5 mm of the same material. A gradient from 1% acetonitrile to 99% acetonitrile in 18 min was applied at 0.4 mL min^-1^ (solvent A 0.1% formic acid in water, solvent B 0.1% formic acid in acetonitrile), before returning to initial conditions. Eluting compounds was sprayed in both positive and negative ion mode by an Apollo II ion funnel ESI source (Bruker Daltonics Inc). Source settings were: capillary voltage 4500 V (positive mode) or 3600 V (negative mode); end plate offset 500 V; drying temperature 220°C; desolvation gas (nitrogen) flow 8.0 l min^-1^; nebulizer gas pressure 2.2 bar. Samples were analysed in timsoff mode, auto MS/MS settings were: switching threshold 500 cts; cycle time 0.5 sec; active exclusion after 3 spectra; release after 0.2 min. Analytes selected for fragmentation were fragmented by collision with nitrogen gas at a collision energy from 20-30 eV depending on precursor m/z. Precursors and fragments were analysed by the time of flight analyser using a range of 20-1300 *m/z*.

### Metabolomics data processing

LC-QTOF-MS data sets from both positive and negative ionization mode were processed using a T-ReX 3D workflow MetaboScape® (v5.0, Bruker) with default settings, but with small modification in peak detection parameters: intensity threshold = 1,000, minimum peak length = 8, retention time range = 0.3 – 20 min, and mass range = 20 – 1200 m/z. Automatic mass calibration was employed using sodium formate, and to correct for batch effects, QC samples were used. Buckets (metabolic features) that were not present in all replicates (n = 5) of each treatment were filtered out. Finally, the data sets from positive (total 536 features) and negative (total 377 features) mode were merged resulting in total 910 features that included 3 merged features and 907 non-merged features. The previous two steps were also done in MetaboScape® as well. Tentative annotation of features was performed using several built-in annotation approaches in MetaboScape®. These were i) searches of several in-house and public libraries of fragment spectra, ii) precursor isotope pattern based calculation of molecular formula, iii) matching of accurate mass, retention time and fragment mass spectra to an in-house built list verified compound standards, iv) searching for candidates based on molecular formula in online compound databases such as Pubchem, Chebi and Chemspider followed by in silico fragmentation analysis of candidate compounds using a built in implementation of the metFrag algorithm (metfrag or MF in results; Ruttkies et al. 2016). Thresholds set for mass accuracy of measured versus theoretical mass (m/z) where: high threshold < 3.0 low threshold < 5.0 ppm. Retention time deviation from retention times of verified compound standards: high threshold < 0.17 low threshold < 0.34 min. Accuracy of isotope modelling accuracy (mSigma value) high threshold < 10 low threshold < 20. MS/MS fragmentation match score: high threshold > 900 low threshold > 600).

Metabolite annotation and structural elucidation were performed using SIRIUS 6.1 (Ludwig et al. 2020a) based on tandem mass spectrometry (MS/MS) data acquired from tomato recombinant inbred line (RIL) root exudates. Peak lists were imported in .*mgf* format and processed through the command-line interface to ensure reproducibility and scalability across multiple samples. The output files were adjusted for high-confidence formula and structure prediction.

Molecular formula determination employed isotope pattern filtering and elemental constraints restricted to the common bioelements, while additional elements (B, Cl, Br, Se, S) were permitted to capture halogenated and sulfur-containing metabolites. Searches were conducted against multiple integrated databases, including BIO, METACYC, CHEBI, COCONUT, GNPS, HMDB, KEGG, PLANTCYC, PUBCHEM, and YMDB, covering a wide spectrum of biologically relevant and chemically diverse compounds. The mass deviation for MS² matching was set to 10 ppm, and heuristic evaluations were applied selectively to ions with m/z below 650 to improve computational efficiency without compromising identification accuracy.

Confidence levels for molecular formula assignment were refined using the ZODIAC algorithm (Ludwig et al. 2020b) through a two-step Bayesian network approach. The model employed 20,000 iterations, a burn-in period of 2000, and 10 Markov chains to assess compositional plausibility based on network connectivity thresholds (--ZodiacEdgeFilterThresholds.thresholdFilter=0.95). A broad range of ion adducts was included ([M+H]+, [M+Na]+, [M–H]–, [M+Cl]–, among others) to account for multiple ionization modes typical of root exudate metabolomes. The final workflow integrated formula, ZODIAC, fingerprint, structure, and CANOPUS modules (Dührkop et al. 2021) to generate comprehensive compound-level annotations and chemical class predictions within a unified analytical framework.

The metabolite feature intensity table obtained from MetaboScape® was imported in R for further analyses. To reduce potential noises from environment, features not significantly different comparing to experimental blank (a pot without plant) were excluded (T-test, *p* < 0.05), and, finally, 690 features were yielded. A pseudonymous name beginning with p was assigned to each unique metabolite feature (e.g. p1, p2, p3, and etc.). Feature intensity table was square-root transformed prior to further analyses. Principal component analysis (PCA) was performed using R basic function ‘prcomp’ with the parameter ‘scale=F’, and permutational multivariate analysis of variance (PERMANOVA) based on Euclidean distance matrix was performed with 1,000 permutations using the function ‘adonis’ in R package vegan (v2.7-2).

In general, data sets were handled using R packages tidyverse (v2.0.0) and the results were visualized using R package ggplot2 (v4.0.0) in R (v4.5.1).

### Heritability and QTL mapping

Phenotypic variance of genotype (*V_G_*) and total population (*V_P_*) were calculated using analysis of variance (ANOVA) to measure broad-sense heritability (*H*^2^ = *V_G_*/*V_P_*) for each feature and its statistical significance (*p* <0.05) was tested using 1,000 times permutation. Quantitative trait locus (QTL) analysis was employed using R package qtl2 (Broman et al. 2019) with previously published genotype probability and physical map of single-nucleotide polymorphism (SNP) markers in RIL population (Sterken et al. 2023). Prior to QTL mapping, genotype (allele AA/BB; AA for MM and BB for SP) was inferred according to the genotype probability score (AA < 0.5 ≤ BB), and median value of metabolite feature intensity was acquired. First, pseudomarkers were inserted into the physical map between SNP markers with a step distance of 1Mb to increase the map resolution, and then new genotype probability for the SNP markers and pseudomarkers was calculated with an error probability of 0.001. With the new genotype probability and median values of feature intensity, a genome scan was employed to find QTL using a single-QTL model. A threshold of logarithm of odd (LOD) score for each QTL was determined at alpha level of 5% using a permutation test (1,000 times).

Annotation and location of genes identified around QTL regions were searched based on *S. lycopersicum* Heinz 1706 genome assembly and annotation SL4.0 and ITAG4.0 using python in-house script and visualized using Jbrowser available at Sol Genomics Network (https://solgenomics.net/).

### Data availability

The raw metabolomics data have been submitted to public data repository Zenodo (DOI: 10.5281/zenodo.17913640). All data sets, scripts and information of packages for the data analysis performed in this study can be found in “https://github.com/BoraKim2018/2025_RIL_tomato_rootexudate”.

## Results

### Composition of the root exudate

LC-QTOF-MS analysis of the chemical composition of the root exudate (RE) of the RIL population showed the presence of a total of 690 metabolic features after data preprocessing (Table. S1). The unique metabolic features were designated with the prefix “p” (e.g. p1, p2, p3…), and this annotation was used throughout this study. The molecular formulae, putative compound names, and compound classes were predicted using Metaboscape and SIRIUS. PCA on the RE features of the RIL population showed that the parents of the population cluster separately along PC1 implying they have a distinctive RE composition (Fig. 1). The individual lines of the population are located around the two parents, and display transgression, with many lines having a RE composition outside the metabolic space defined by the two parents. The individual replicates of the parents do not cluster strongly together, indicating variation among the biological replicates, which is inherent to the methods used. This is reflected in the large residual variation found in PERMANOVA based on Euclidean distance of the RE profile of the two parents (Table 1). Nevertheless, the analysis showed that 23% of the total variance was explained by genotype (*p* <0.05; Table. 1).

**Figure 1.**
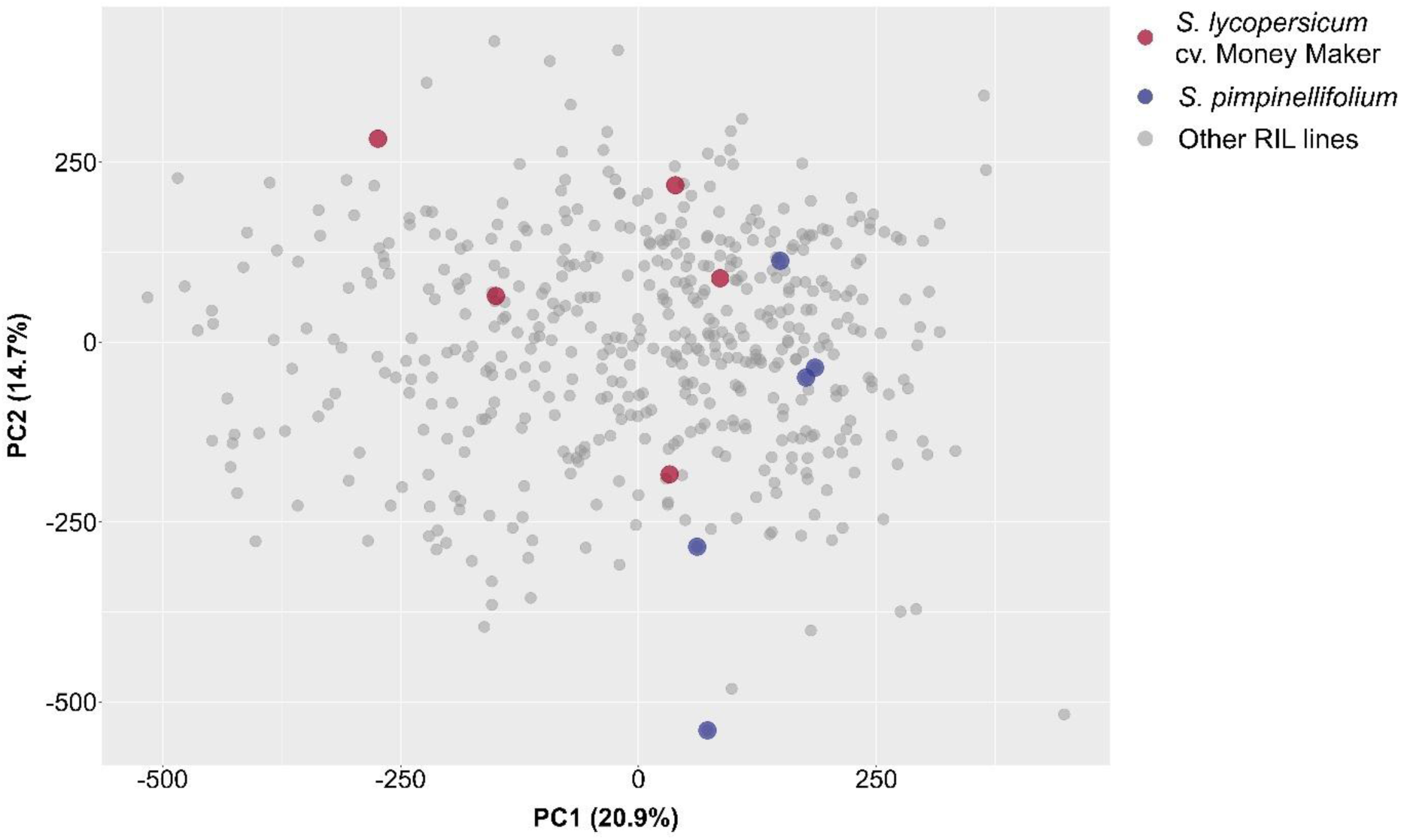
Principal component analysis on the metabolite features detected in root exudate obtained from the tomato recombinant inbred line (RIL) and the two parents. Each dot represents a sample point.

**Table 1.** Result of permutational multivariate analysis of variance (PERMANOVA) based on Euclidean distance of root exudate composition obtained from *Solanum lycopersicum* cv. Moneymaker and *S. pimpinellifolium*. PERMANOVA was performed with 1,000 permutations.

|  | Degrees of freedom | Sums of squares | Mean square | F Model | R<sup>2</sup> | Pr (>F) |
| --- | --- | --- | --- | --- | --- | --- |
| Genotype | 1 | 384042 | 384042 | 02.4086 | 0.23141 | 0.033 |
| Residuals | 8 | 1275558 | 159445 |  | 0.76859 |  |
| Total | 9 | 1659600 |  |  | 1.00000 |  |

### Heritability

To investigate how genetic variation in the RIL population affects the composition of the RE, broad-sense heritability (*H*^2^) for each phenotypic trait (i.e. metabolic feature) was analyzed. A total of 53 features displayed a significant *H*^2^ (Fig. 2). The majority of these metabolic features had an *H*^2^ within the range of 0.23 - 0.49, except for p576 (predicted as dihydroactinidiolide) displaying the highest *H*^2^ of 8.42.

**Figure 2.**
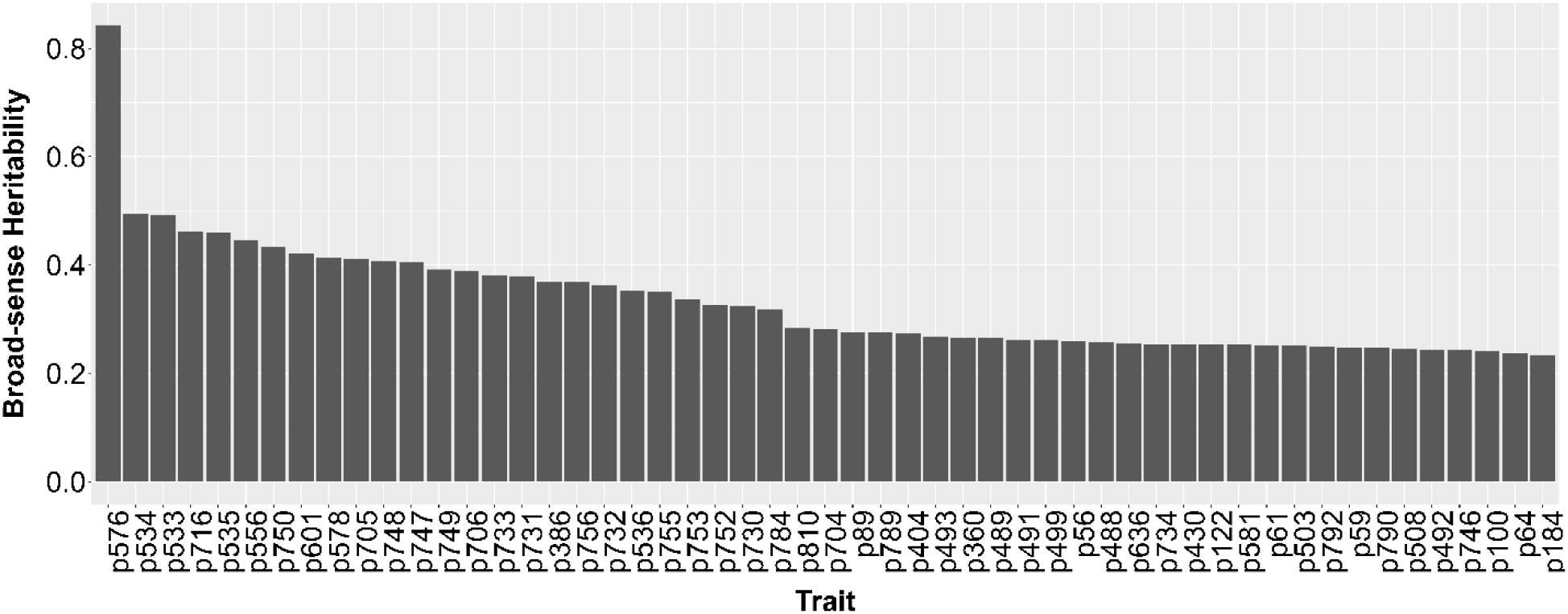
Broad-sense heritability (H²) of metabolite features (labeled with prefix “p”) detected in root exudates from tomato recombinant inbred lines. H² was estimated using analysis of variance to partition variance attributable to genotype versus total phenotypic variance. Statistical significance (p < 0.05) was determined by 1,000 permutations of the variance components. Only features with significant H² are shown.

### Quantitative trait locus analysis

Quantitative trait locus (QTL) analysis to find tomato genomic regions associated with metabolic features in the RE of the RIL population, yielded a total of 82 QTLs, of which 69 were unique SNPs, with an association with 7 metabolic features (Fig. 3; Table 2). Among the 82 QTLs, 55 were mapped with allele AA from parent MM, and 27 were mapped with allele BB from the other parent SP. Chromosome (Ch) 4 harbored the biggest number of QTLs (11), followed by Ch 1 (10 QTLs) and Ch 8 (9 QTLs). A list of genes within each QTL region was retrieved using the *S. lycopersicum* Heinz 1706 reference genome and gene model annotation SL4.0 (Table S2).

**Figure 3.**
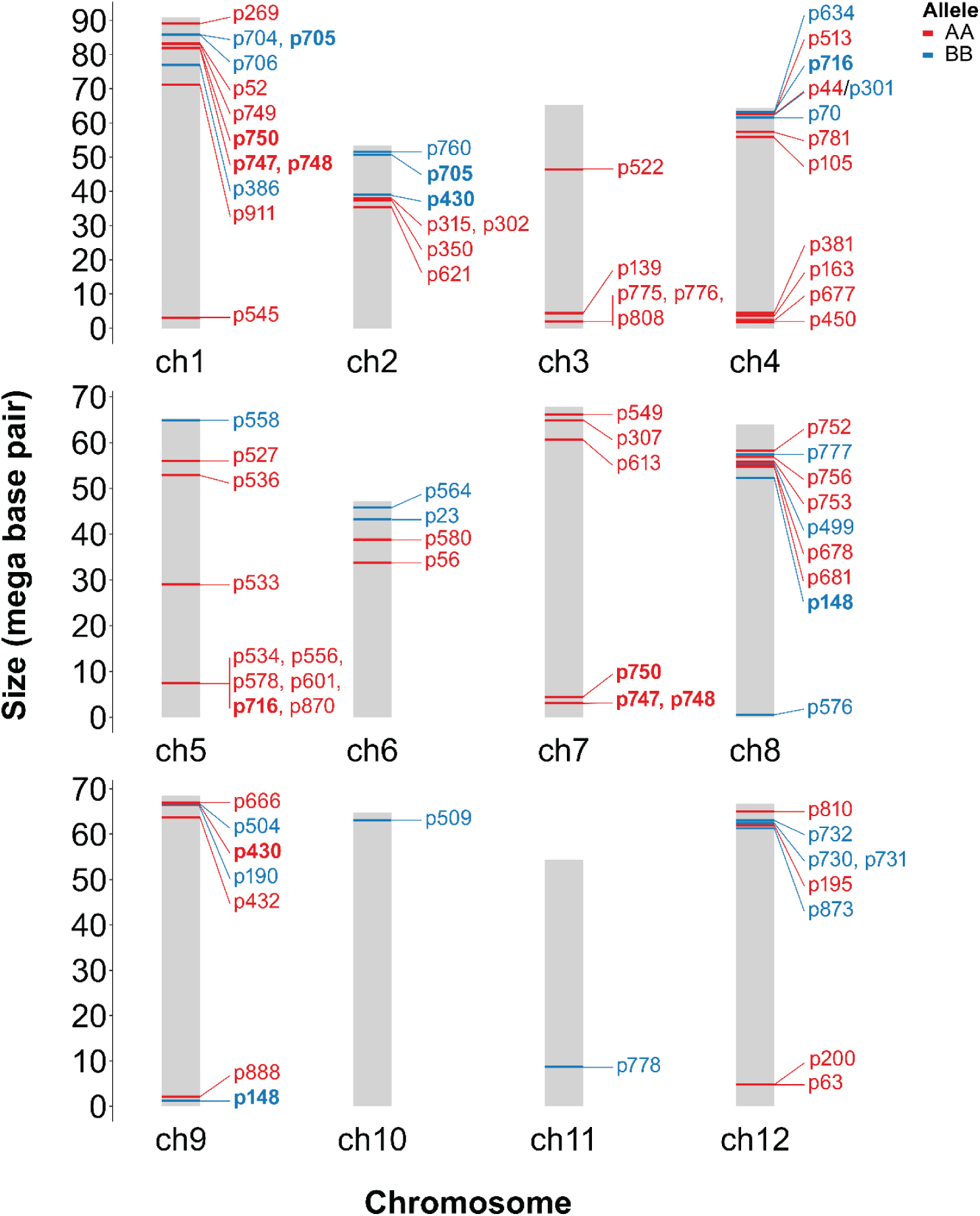
Genomic locations of single nucleotide polymorphisms (SNPs) significantly associated with metabolic features (labeled with prefix “p”) detected in the root exudate of tomato recombinant inbred lines. SNP alleles are annotated as originating from *Solanum lycopersicum* cv. Moneymaker (AA) or *S. pimpinellifolium* (BB), indicating parental allele contributions to metabolite variation. Bold font indicates metabolic features that associate with multiple SNPs.

**Table 2.** Quantitative trait loci associated with metabolic features in tomato root exudate.

| Trait | LOD<sup>a</sup> | SNP<sup>b</sup> | Ch<sup>c</sup> | Pos<sup>d</sup><br>(Mb) | Interval | Allele | RT<sup>e</sup> | m/z | ESI<sup>f</sup> | Molecular formula<br>(annotation)<sup>g</sup> | Predicted<br>subclass/class<br>(probability 0-1)<sup>h</sup> |
| --- | --- | --- | --- | --- | --- | --- | --- | --- | --- | --- | --- |
| p545 | 4.010 | snp97 | 1 | 3.147 | 2.502 -<br>4.157 | AA | 7.900 | 417.192 | [M-H]- | C15H36N2O7P2 | Tetraketide<br>meroterpenoids<br>(0.692) |
| p911 | 3.775 | snp208 | 1 | 71.240 | 0.506 -<br>76.252 | AA | 19.890 | 74.937 | [M+H]+ |  |  |
| p386 | 5.819 | snp312 | 1 | 76.974 | 76.387 -<br>78.552 | BB | 5.830 | 295.102 | [M+H]+ | C11H18O9 | Disaccharides (1) |
| p748 | 4.656 | snp376 | 1 | 81.947 | 80.722 -<br>86.827 | AA | 11.410 | 247.117 | [M+H]+ | C11H18O6 |  |
| p747 | 4.205 | snp376 | 1 | 81.947 | 80.722 -<br>86.859 | AA | 11.410 | 85.065 | [M+H]+ | C5H8O (3-Methylcrotonaldeh<br>yde) | Monoterpenoids<br>(0.756) |
| p750 | 4.341 | snp378 | 1 | 81.990 | 80.898 -<br>86.827 | AA | 11.410 | 181.086 | [M+H]+ | C10H12O3 | Cyclic<br>polyketides<br>(0.169) |
| p749 | 4.099 | snp384 | 1 | 83.064 | 80.898 -<br>86.55 | AA | 11.410 | 57.070 | [M+H]+ |  |  |
| p52 | 3.490 | snp389 | 1 | 83.402 | 0.01 -<br>89.626 | AA | 0.530 | 113.989 | [M-H]- |  |  |
| p706 | 6.121 | snp422 | 1 | 85.842 | 85.251 -<br>87.274 | BB | 10.480 | 57.070 | [M+H]+ |  |  |
| p705 | 7.399 | snp423 | 1 | 85.847 | 85.56 -<br>86.55 | BB | 10.480 | 247.118 | [M+H]+ | C11H18O6 |  |
| p704 | 3.914 | snp423 | 1 | 85.847 | 80.109 -<br>87.57 | BB | 10.480 | 116.928 | [M-H]- |  | Fatty acyl<br>glycosides of<br>mono- and<br>disaccharides<br>(0.756) |
| p269 | 3.238 | snp486 | 1 | 89.073 | 0.01 -<br>90.864 | AA | 3.520 | 246.055 | [M+H]+ |  | Lysine alkaloids<br>(0.237) |
| p621 | 3.320 | snp567 | 2 | 35.425 | 13.494 -<br>43.34 | AA | 9.050 | 401.088 | [M-H]- | C15H15N8O4P | Simple phenolic<br>acids (0.408) |
| p350 | 3.604 | snp597 | 2 | 37.411 | 16.713 -<br>43.34 | AA | 4.960 | 397.246 | [M-H]- | C13H35N8O4P | Small peptides<br>(0.694) |
| p315 | 4.307 | snp613 | 2 | 38.045 | 36.878 -<br>39.056 | AA | 4.370 | 415.256 | [M-H]- | C13H37N8O5P | Dipeptides<br>(0.892) |
| p302 | 3.437 | snp613 | 2 | 38.045 | 35.153 -<br>39.637 | AA | 4.080 | 259.165 | [M+H]+ | C12H22N2O4 | Small peptides<br>(0.764) |
| p430 | 3.467 | snp631 | 2 | 39.062 | 16.713 -<br>51.96 | BB | 6.590 | 331.255 | [M+H+<br>H]<sub>2</sub><sup>+</sup> | C35H68N2O9 |  |
| p705 | 3.984 | snp862 | 2 | 50.722 | 46.235 -<br>53.473 | BB | 10.480 | 247.118 | [M+H]+ | C11H18O6 | Fatty acyl<br>glycosides of<br>mono- and<br>disaccharides<br>(0.756) |
| p760 | 3.325 | snp880 | 2 | 51.574 | 45.343 -<br>53.473 | BB | 11.600 | 106.086 | [M+H]+ |  |  |
| p775 | 4.003 | snp978 | 3 | 2.053 | 0 -<br>65.298 | AA | 12.280 | 277.237 | [M+H]+ | C15H32O4 | Monoacylglycerol<br>s (0.89) |
| p808 | 3.865 | snp978 | 3 | 2.053 | 0 -<br>65.298 | AA | 13.250 | 291.253 | [M+H]+ | C16H34O4 | Monoacylglycerol<br>s (0.886) |
| p776 | 3.549 | snp978 | 3 | 2.053 | 0 -<br>65.298 | AA | 12.280 | 299.219 | [M+H]+ | C13H26N6O2 | Fatty esters<br>(0.855) |
| p139 | 3.447 | snp1044 | 3 | 4.454 | 3.715 -<br>47.744 | AA | 0.800 | 180.973 | [M-H]- | C4H7O4PS | Dicarboxylic<br>acids (0.481) |
| p522 | 3.309 | snp1101 | 3 | 46.521 | 6.645 -<br>46.593 | AA | 7.680 | 411.167 | [M-H]- | C12H34N2O9P2 | Tryptophan<br>alkaloids (0.609) |
| p450 | 3.427 | snp1403 | 4 | 1.858 | 0 -<br>58.72 | AA | 6.700 | 117.929 | [M-H]- |  |  |
| p677 | 3.432 | snp1427 | 4 | 2.526 | 0.49 -<br>63.868 | AA | 9.930 | 281.140 | [M-H]- | C15H22O5 (5,9-dihydroxy-7-<br>(hydroxymethyl)-<br>5,7-dimethyl-4,5a,6,8,8a,9-hexahydro-1H-azuleno[5,6-c]furan-3-one) |  |
| p163 | 3.307 | snp1458 | 4 | 3.642 | 2.758 -<br>51.194 | AA | 1.280 | 195.811 | [M-H]- |  |  |
| p381 | 3.510 | snp1474 | 4 | 4.470 | 2.062 -<br>53.946 | AA | 5.710 | 497.334 | [M-H]- | C25H46N4O6 | Steroids<br>(0.512) |
| p105 | 3.754 | snp1585 | 4 | 55.989 | 3.236 -<br>56.744 | AA | 0.630 | 564.830 | [M-H]- |  | Flavonoids<br>(0.378) |
| p781 | 3.514 | snp1608 | 4 | 57.450 | 0.508 -<br>59.69 | AA | 12.420 | 303.231 | [M+H]+ | C20H30O2 (8-[5]-<br>ladderane-octanoic<br>acid) | Diterpenoids<br>(0.976) |
| p70 | 5.133 | snp1715 | 4 | 61.613 | 61.286 -<br>63.098 | BB | 0.560 | 378.918 | [M-H]- |  | Pseudoalkaloids<br>(0.431) |
| p44 | 3.477 | snp1724 | 4 | 62.615 | 4.123 -<br>64.46 | AA | 0.520 | 453.926 | [M-H]- | C10H9N5O8S4 | Pyrrole alkaloids<br>(0.271) |
| p301 | 3.354 | snp1724 | 4 | 62.615 | 2.341 -<br>64.46 | BB | 4.070 | 229.105 | [M+H]+ | C7H12N6O3 | Fatty Acids and<br>Conjugates<br>(0.129) |
| p716 | 3.490 | snp1731 | 4 | 62.753 | 0.344 -<br>64.46 | BB | 10.650 | 261.133 | [M+H]+ |  | Macrolide<br>lactones (0.637) |
| p513 | 3.980 | snp1739 | 4 | 62.935 | 62.615 -<br>63.811 | AA | 7.620 | 54.041 | [M+H]+ |  |  |
| p634 | 3.288 | snp1743 | 4 | 63.245 | 0 -<br>64.46 | BB | 9.290 | 561.270 | [M-H]- |  | Triterpenoids<br>(0.554) |
| p534 | 4.401 | snp1915 | 5 | 7.419 | 7.41 -<br>55.99 | AA | 7.750 | 565.251 | [M-H]- | C21H43N8O4PS2 |  |
| p601 | 3.820 | snp1915 | 5 | 7.419 | 7.208 -<br>58.514 | AA | 8.700 | 579.267 | [M-H]- | C20H48N6O7S3 |  |
| p578 | 6.182 | snp1915 | 5 | 7.419 | 7.41 -<br>58.514 | AA | 8.400 | 579.267 | [M-H]- | C20H48N6O7S3 | Iridoids<br>monoterpenoids<br>(0.881) |
| p556 | 3.529 | snp1915 | 5 | 7.419 | 7.41 -<br>55.99 | AA | 8.070 | 566.254 | [M-H]- | C14H42N13O5PS2 | Macrolide<br>lactones (0.637) |
| p716 | 3.276 | snp1915 | 5 | 7.419 | 7.41 -<br>55.99 | AA | 10.650 | 261.133 | [M+H]+ |  | Tripeptides<br>(0.532) |
| p810 | 3.530 | snp1915 | 5 | 7.419 | 6.943 -<br>59.393 | AA | 13.250 | 519.283 | [M-H]- |  | Tryptophan<br>alkaloids (0.124) |
| p533 | 4.319 | cch05.loc2<br>9 | 5 | 29.000 | 7.41 -<br>55.99 | AA | 7.750 | 611.256 | [M-H]- | C26H44O16 | Sesquiterpenoids<br>(0.565) |
| p536 | 4.116 | snp1927 | 5 | 52.882 | 7.41 -<br>58.482 | AA | 7.750 | 81.033 | [M+H]+ | C5H4O | Cyclic peptides<br>(0.508) |
| p527 | 4.464 | snp1929 | 5 | 55.990 | 7.576 -<br>58.61 | AA | 7.740 | 116.928 | [M-H]- |  | Polyols (0.183) |
| p558 | 3.372 | snp2103 | 5 | 64.871 | 3.118 -<br>65.269 | BB | 8.080 | 387.181 | [M-H]- | C14H34N2O6P2 | Small peptides<br>(0.372) |
| p56 | 4.201 | snp2179 | 6 | 33.809 | 2.517 -<br>42.919 | AA | 0.540 | 386.925 | [M+H]+ |  |  |
| p580 | 3.401 | snp2287 | 6 | 38.807 | 35.297 -<br>39.3 | AA | 8.440 | 100.933 | [M-H]- |  |  |
| p23 | 3.423 | snp2379 | 6 | 43.207 | 41.536 -<br>44.802 | BB | 0.370 | 261.851 | [M-H]- |  |  |
| p564 | 3.895 | snp2415 | 6 | 45.834 | 40.943 -<br>46.949 | BB | 8.160 | 473.182 | [M-H]- | C20H27N8O4P | Steroids<br>(0.706) |
| p748 | 2.947 | snp2524 | 7 | 3.093 | 2.377 -<br>59.41 | AA | 11.410 | 247.117 | [M+H]+ | C11H18O6 |  |
| p747 | 2.919 | snp2524 | 7 | 3.093 | 2.271 -<br>57.87 | AA | 11.410 | 85.065 | [M+H]+ | C5H8O (3-Methylcrotonaldeh<br>yde) | Monoterpenoids<br>(0.756) |
| p750 | 2.995 | snp2573 | 7 | 4.404 | 2.377 -<br>59.41 | AA | 11.410 | 181.086 | [M+H]+ | C10H12O3 | Cyclic<br>polyketides<br>(0.169) |
| p613 | 3.741 | snp2648 | 7 | 60.626 | 59.294 -<br>65.393 | AA | 8.860 | 135.080 | [M+H]+ | C9H10O<br>(Chavicol) | Cinnamic acids<br>and derivatives<br>(0.741) |
| p307 | 4.180 | snp2718 | 7 | 64.905 | 62.684 -<br>65.529 | AA | 4.260 | 257.099 | [M+H]+ | C8H12N6O4 | Phenolic acids<br>(C6-C1) (0.775) |
| p549 | 3.878 | snp2743 | 7 | 66.180 | 63.055 -<br>67.884 | AA | 7.990 | 216.196 | [M+H]+ | C12H25NO2 | Fatty amides<br>(0.851) |
| p576 | 33.575 | snp2770 | 8 | 0.575 | 0.575 -<br>0.582 | BB | 8.370 | 181.122 | [M+H]+ | C11H16O2<br>(Dihydroactinidioli<br>de) | Monoterpenoids<br>(0.709) |
| p148 | 4.097 | snp2939 | 8 | 52.302 | 13.505 -<br>55.823 | BB | 0.960 | 173.871 | [M-H]- |  |  |
| p681 | 3.609 | snp2983 | 8 | 54.751 | 4.472 -<br>56.88 | AA | 10.010 | 121.028 | [M+H]+ | C7H4O2 | Coumarins<br>(0.322) |
| p678 | 3.656 | snp2986 | 8 | 54.997 | 2.028 -<br>57.046 | AA | 10.010 | 65.038 | [M+H]+ |  | Phenylpropanoids<br>(C6-C3) (0.215) |
| p499 | 3.636 | snp2999 | 8 | 55.569 | 1.269 -<br>57.479 | BB | 7.460 | 407.171 | [M-H]- | C15H29N4O7P | Quassinoids<br>(0.695) |
| p753 | 3.984 | snp3001 | 8 | 55.814 | 13.505 -<br>58.448 | AA | 11.510 | 187.060 | [M+H]+ | C8H10O5 | Cyclic<br>polyketides<br>(0.876) |
| p756 | 3.621 | snp3016 | 8 | 56.873 | 55.126 - 58.118 | AA | 11.520 | 317.159 | [M+H]<sup>+</sup> | C15H24O7 | Open-chain polyketides (0.468) |
| p777 | 5.458 | snp3020 | 8 | 57.471 | 56.041 - 57.773 | BB | 12.290 | 61.989 | [M-H]<sup>-</sup> |  |  |
| p752 | 3.525 | snp3038 | 8 | 58.204 | 0.589 - 59.035 | AA | 11.500 | 127.039 | [M+H]<sup>+</sup> | C6H6O3 | Phenolic acids (C6-C1) (0.737) |
| p148 | 3.317 | snp3180 | 9 | 1.232 | 0 - 68.514 | BB | 0.960 | 173.871 | [M-H]<sup>-</sup> |  |  |
| p888 | 3.408 | snp3210 | 9 | 2.127 | 0 - 68.514 | AA | 18.300 | 145.924 | [M-H]<sup>-</sup> |  |  |
| p432 | 3.531 | snp3455 | 9 | 63.659 | 26.491 - 68.514 | AA | 6.590 | 276.728 | [M+H+H]<sup>2+</sup> | C30H57N5O4 |  |
| p190 | 3.723 | snp3541 | 9 | 66.437 | 1.771 - 68.514 | BB | 2.020 | 73.945 | [M+H]<sup>+</sup> |  |  |
| p430 | 3.975 | snp3543 | 9 | 66.448 | 63.179 - 67.188 | AA | 6.590 | 331.255 | [M+H+H]<sup>2+</sup> | C35H68N2O9 |  |
| p504 | 4.632 | snp3556 | 9 | 66.875 | 66.437 - 68.514 | BB | 7.540 | 381.156 | [M-H]<sup>-</sup> | C12H34N2O5S3 | Nucleosides (0.784) |
| p666 | 3.760 | cch09.loc67 | 9 | 67.000 | 14.301 - 68.514 | AA | 9.690 | 547.290 | [M-H]<sup>-</sup> |  | Triterpenoids (0.734) |
| p509 | 4.029 | snp3901 | 10 | 63.111 | 61.985 - 63.703 | BB | 7.600 | 449.218 | [M-H]<sup>-</sup> | C16H40N2O8P2 | Cardenolides (0.861) |
| p778 | 3.708 | snp4070 | 11 | 8.674 | 6.386 - 47.568 | BB | 12.340 | 321.263 | [M+H]<sup>+</sup> | C17H36O5 | Monoacylglycerols (0.647) |
| p63 | 3.443 | snp4348 | 12 | 4.783 | 2.398 - 61.941 | AA | 0.550 | 317.950 | [M-H]<sup>-</sup> | C9H9N3O2S4 | Small peptides (0.514) |
| p200 | 5.143 | snp4352 | 12 | 4.800 | 4.033 - 57.07 | AA | 2.140 | 239.903 | [M-H]<sup>-</sup> |  |  |
| p873 | 4.243 | snp4386 | 12 | 61.349 | 5.178 - 61.526 | BB | 16.190 | 57.070 | [M+H]<sup>+</sup> | C4H8 |  |
| p195 | 4.047 | snp4400 | 12 | 61.962 | 3.516 - 63.426 | AA | 2.080 | 143.935 | [M-H]<sup>-</sup> |  |  |
| p731 | 4.122 | snp4414 | 12 | 62.461 | 3.516 - 65.68 | BB | 10.970 | 187.060 | [M+H]<sup>+</sup> | C8H10O5 |  |
| p730 | 3.654 | snp4414 | 12 | 62.461 | 3.516 - 66.688 | BB | 10.970 | 71.085 | [M+H]<sup>+</sup> |  | Phenolic acids (0.343) |
| p732 | 4.293 | snp4431 | 12 | 63.005 | 5.406 - 65.592 | BB | 10.970 | 127.039 | [M+H]<sup>+</sup>, [M+H-H2O]<sup>+</sup> | C6H6O3 | 4-pyrone derivatives (0.627) |
| p810 | 4.137 | snp4490 | 12 | 65.000 | 64.775 - 66.243 | AA | 13.250 | 519.283 | [M-H]<sup>-</sup> |  | Sesquiterpenoids (0.565) |

<sup>a</sup>LOD, logarithm of odds; <sup>b</sup>SNP, single nucleotide polymorphism; <sup>c</sup>Ch, Chromosome; <sup>d</sup>Pos, position of SNP; <sup>e</sup>RT, retention time; <sup>f</sup>ESI, electrospray ionization; <sup>g</sup>Molecular formula (annotation), based on in-house and public libraries (see more details in Materials and Methods); <sup>h</sup>Predicted subclass/class (probability 0-1), using SIRIUS

While most QTLs exhibited LOD scores ranging from 2.919 to 7.398, the QTL associated with metabolite feature p576—annotated as dihydroactinidiolide—stood out with an exceptionally high LOD score of 33.575, located at 0.575 Mb on Ch 8 (Table 2; Fig. 4A), consistent with its high heritability (Fig. 2). The median intensity of p576 across genotypes revealed a near-binary distribution, with plants carrying the BB allele exhibiting markedly higher levels than those with the AA allele (Fig. 4B). Notably, this QTL spans an especially narrow genomic interval (0.575–0.582 Mb), containing only three annotated genes: unknown protein (Solyc08g005700.1), and two terpene synthases (Solyc08g005705.1 and Solyc08g005710.3; Fig 4C; Table S2). Interestingly, the surrounding genomic region is enriched in additional terpene synthases (Solyc08g005640.4.1, Solyc08g005677.1.1, and Solyc08g005720.4.1), and an ABC transporter G family member (Solyc08g005580.3.1).

**Figure 4.**
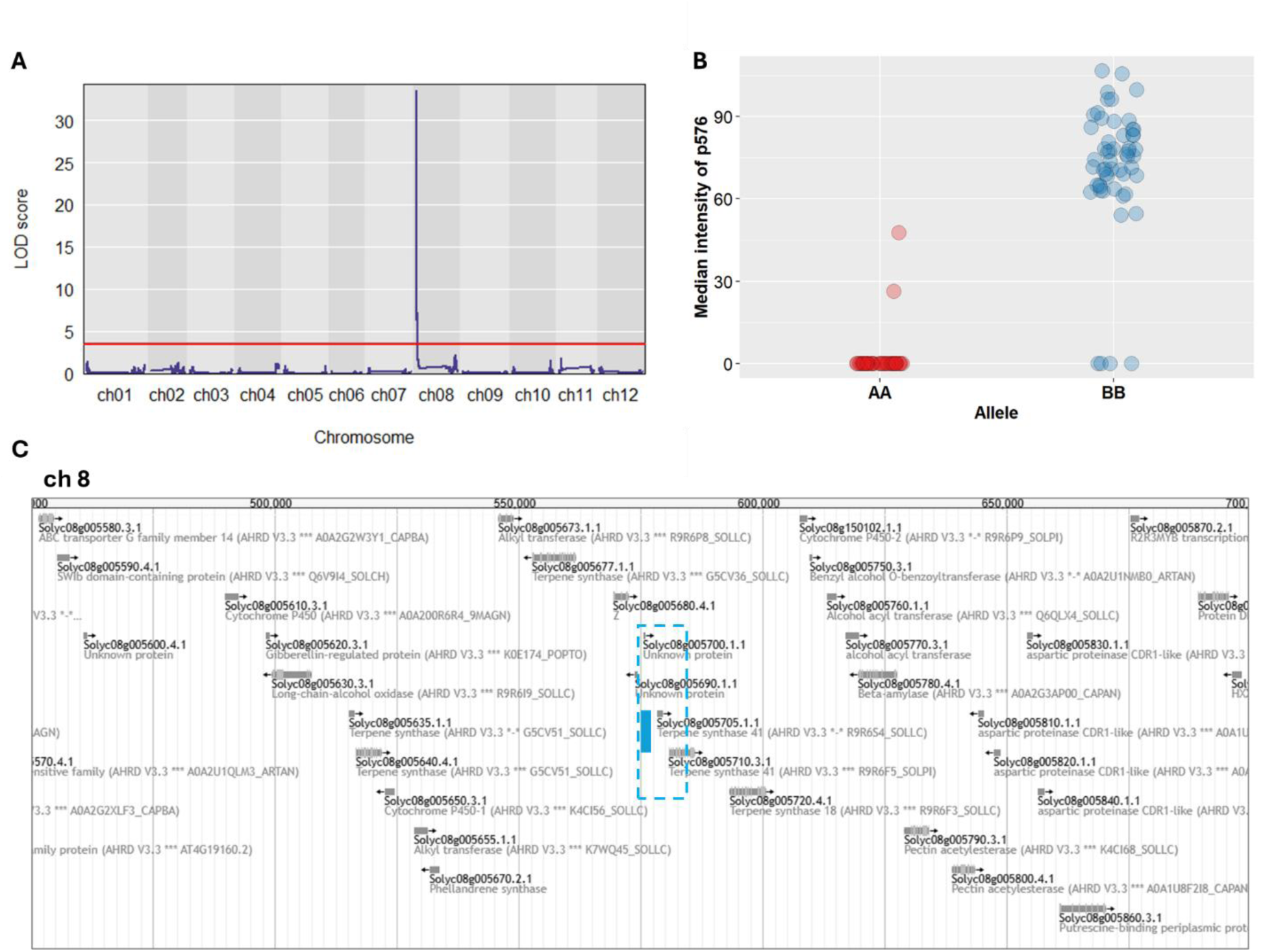
QTL associated with metabolic feature p576. (A) Genome-wide scan showing the logarithm of odds (LOD) score for the QTL. (B) Median intensity of p576 at the SNP marker located at 0.575 Mb on chromosome (ch) 8. Alleles are denoted as originating from *Solanum lycopersicum* cv. Moneymaker (AA) or *S. pimpinellifolium* (BB), illustrating parental allele contributions to metabolite variation. (C) Genomic region surrounding the SNP marker (highlighted with a blue line) within the QTL interval (box indicated with blue dotted line) visualized with JBrowse at solgenomics.net, based on gene model ITAG4.0.

Some metabolic features with significant QTL association could only be putatively identified. For example, p747 was predicted to be 3-methylcrotonaldehyde, which is an oxidized derivative of cytokinin, and showed higher intensity in root exudates of genotypes carrying the AA allele (Fig. 5B). This metabolic feature was associated with two SNP markers located at 81.947 Mb on Ch 1 and 3.093 Mb on Ch 7 (Table 2; Fig. 5A). The corresponding QTL intervals spanned from 80.722 to 86.859 Mb on Ch 1 (harboring 802 genes) and from 2.271 to 57.870 Mb on Ch 7 (harboring 1,025 genes; Table S2). Several genes in these intervals and near the SNP markers are potentially involved in plant stress responses and the cytokinin pathway. For example, acyl-CoA-binding domain-containing protein 4 (Solyc01g099350.3.1), calcium-dependent lipid-binding family protein (Solyc01g099370.3.1), and WD-40 repeat containing protein (Solyc01g099400.4.1) in Ch 1 and 4-coumarate:CoA ligase (Solyc07g008360.2.1) and O-fucosyltransferase family protein (Solyc07g008290.4.1) in Ch 7 (Fig 5C; Table S2).

**Figure 5.**
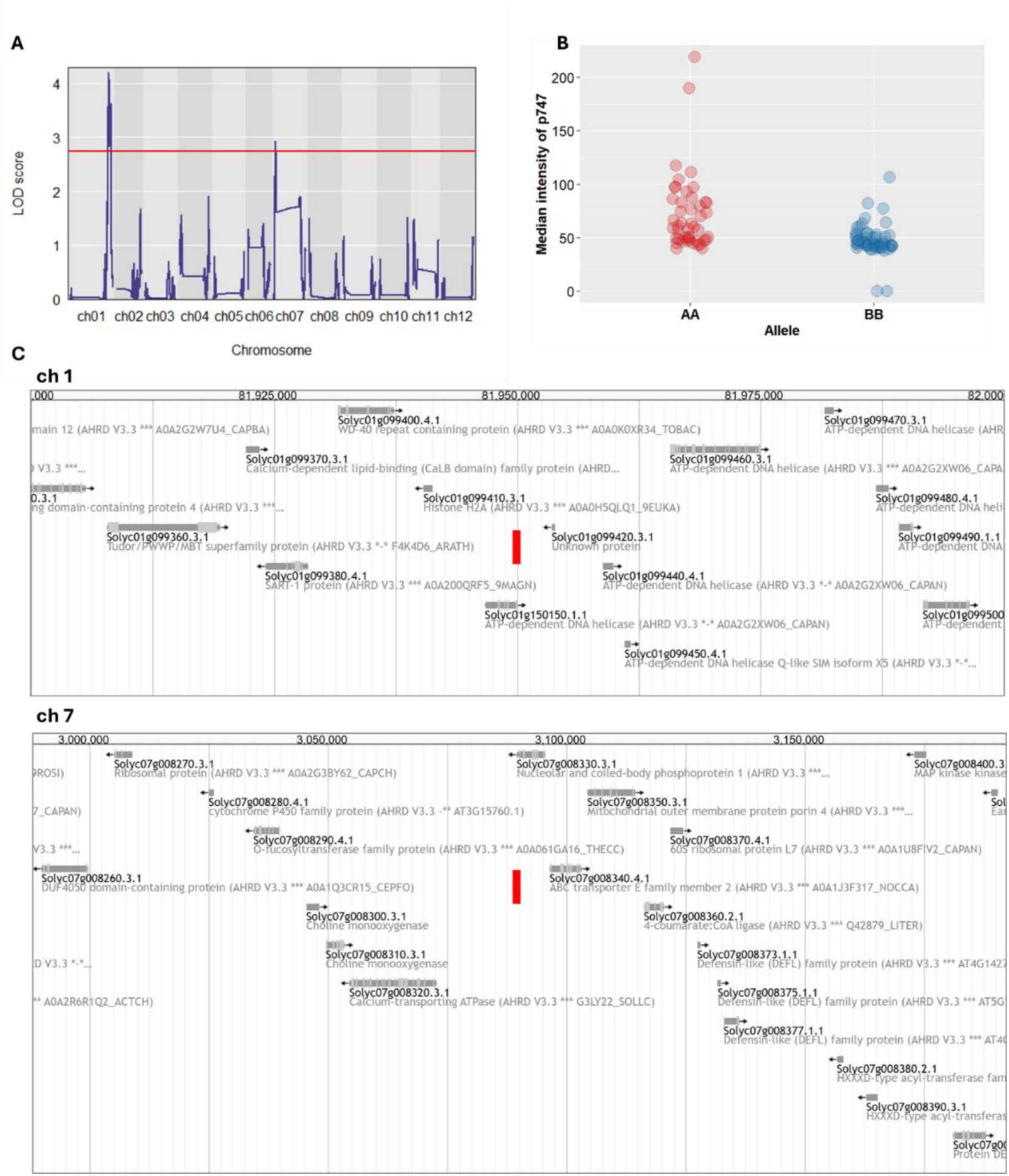
QTL associated with metabolic feature p747. (A) Genome-wide scan showing the logarithm of odds (LOD) scores for the QTLs. (B) Median intensity of p747 at one of the SNP markers located at 81.95 Mb on chromosome (Chr) 1. Alleles are derived from *Solanum lycopersicum* cv. Moneymaker (AA) or *S. pimpinellifolium* (BB), illustrating parental allele contributions to metabolite variation. (C) Genomic region surrounding the SNP markers (highlighted with a red line) visualized using JBrowse at solgenomics.net, based on gene model ITAG4.

Another intriguing metabolic feature, p613—predicted to be chavicol—was associated with a SNP at 60.626 Mb on Ch 7 and mapped to a broad QTL interval spanning 3.740–59.294 Mb, encompassing 713 genes (Table 2; Fig. 6; Table S2). Notably, a gene encoding an S-adenosyl-L-methionine-dependent methyltransferase (Solyc07g052110.4.1) is located in close proximity to the SNP. This enzyme may catalyze the methylation of chavicol, a reaction that has not yet been characterized in tomato (Fig. 6C).

**Figure 6.**
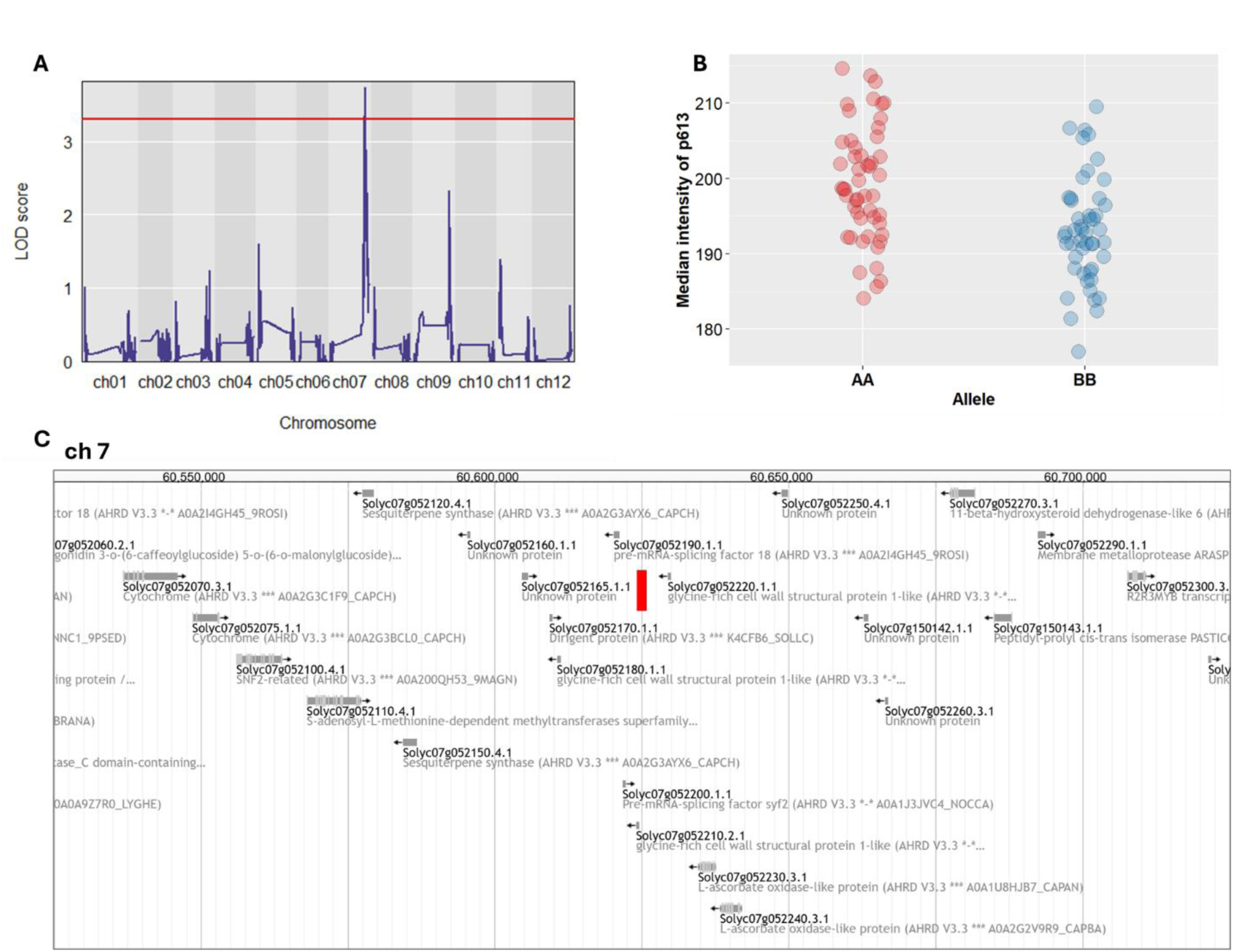
QTL associated with metabolic feature p613. (A) Genome-wide scan showing the logarithm of odds (LOD) scores for the QTL. (B) Median intensity of p613 at the SNP marker located at 60.626 Mb on chromosome (ch) 7. Alleles are denoted as originating from *Solanum lycopersicum* cv. Moneymaker (AA) or *S. pimpinellifolium* (BB), illustrating parental allele contributions to metabolite variation. (C) Genomic region surrounding the SNP marker (highlighted with a red line) visualized with JBrowse at solgenomics.net, based on gene model ITAG4.0.

A number of QTLs located adjacent to genes with potentially important roles in plant metabolism. For instance, metabolic features p666 and p634, predicted to be terpenoids with probability scores of 0.734 and 0.55, respectively. Feature p666 associated with a QTL at 67.000 Mb on Ch 9 positioned directly next to a gene encoding an ATP-binding cassette (ABC) transporter Pleiotropic Drug Resistance (PDR) protein (Solyc09g091660.3.1), which may function in terpenoid transport (Table 2; Fig. 7; Table S2). The other metabolic feature, p634 was associated with a QTL at 63.245 Mb on Ch 4 and the SNP marker is adjacent to several genes involved in auxin signaling (Table2; Fig. 8; Table S2), including Auxin Response Factor 5 (Solyc04g081240.2.1), Small Auxin Up-Regulated RNA 51 (Solyc04g081250.1.1), and Small Auxin Up-Regulated RNA 52 (Solyc04g081270.1.1).

**Figure 7.**
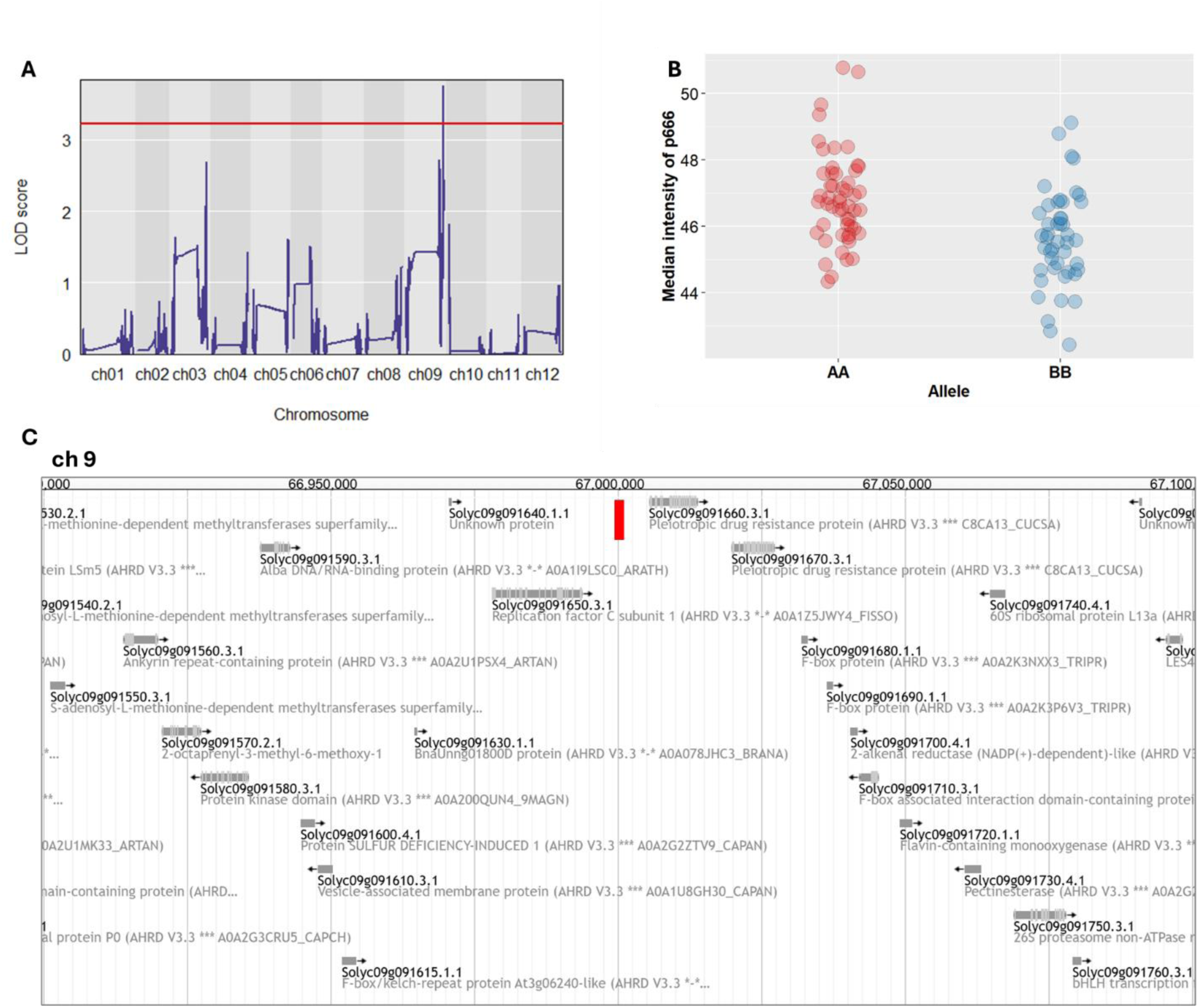
QTL associated with metabolic feature p666. (A) Genome-wide scan showing the logarithm of odds (LOD) scores for the QTL. (B) Median intensity of p666 at the SNP marker located at 67.000 Mb on chromosome (ch) 9. Alleles are denoted as originating from *Solanum lycopersicum* cv. Moneymaker (AA) or *S. pimpinellifolium* (BB), illustrating parental allele contributions to metabolite variation. (C) Genomic region surrounding the SNP marker (highlighted with a red line) visualized with JBrowse at solgenomics.net, based on gene model ITAG4.0.

**Figure 8.**
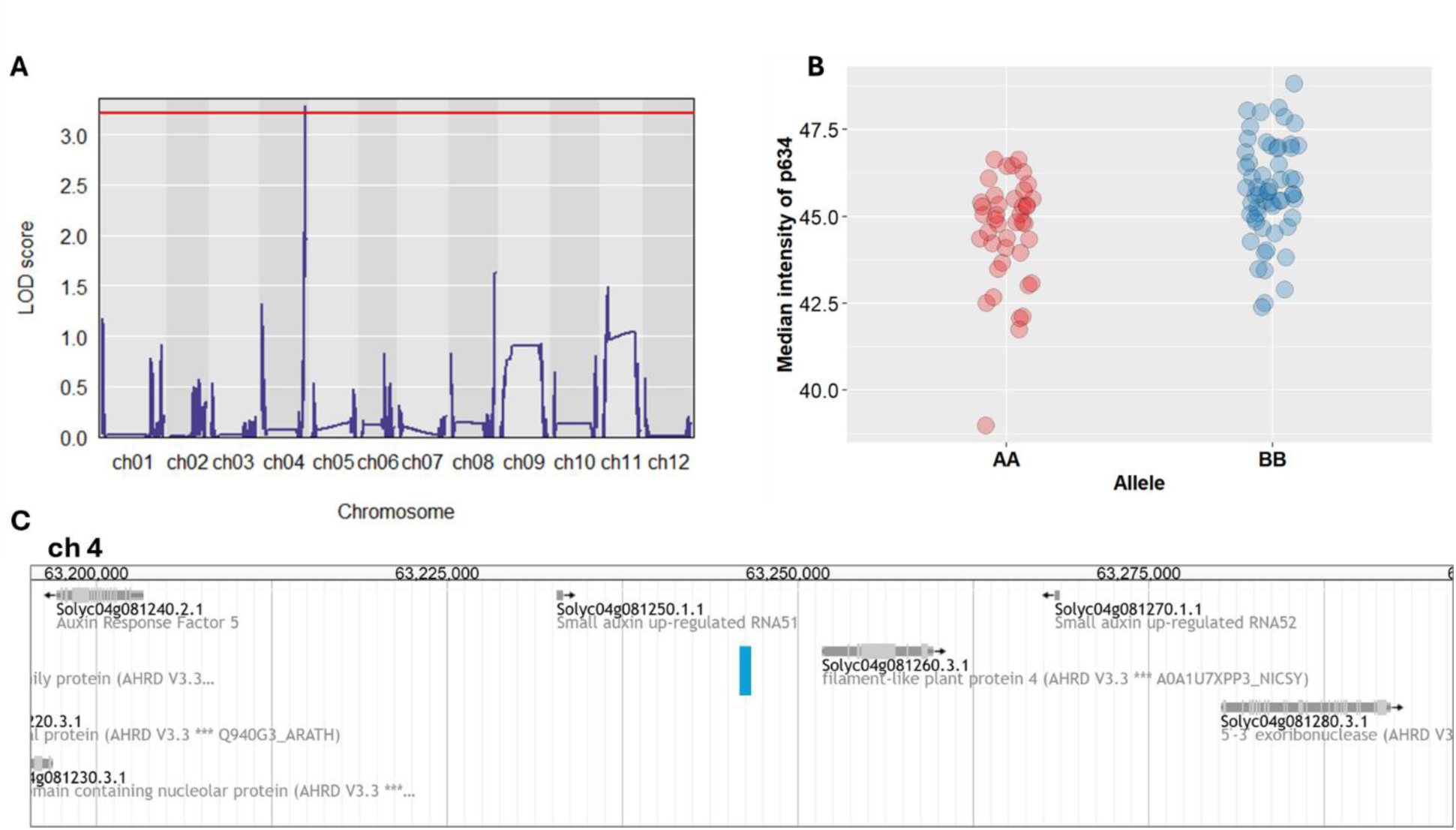
QTL associated with metabolic feature p634. (A) Genome-wide scan showing the logarithm of odds (LOD) scores for the QTL. (B) Median intensity of p634 at the SNP marker located at 63.245 Mb on chromosome (ch) 4. Alleles are denoted as originating from *Solanum lycopersicum* cv. Moneymaker (AA) or *S. pimpinellifolium* (BB), illustrating parental allele contributions to metabolite variation. (C) Genomic region surrounding the SNP marker (highlighted with a blue line) visualized with JBrowse at solgenomics.net, based on gene model ITAG4.0.

Several QTLs were associated with multiple root exudate (RE) metabolic features that appear to be functionally related based on their predicted classifications. For instance, p534, p555, p578, and p601 all contain nitrogen (N) in their predicted molecular formulas and all mapped to a QTL at 7.419 Mb on Ch 5 (Table 2; Fig. 9). Interestingly, the SNP associated with these features was located adjacent to a gene encoding Cysteine Protease 8 (Solyc05g013920.4.1), an enzyme involved in protein degradation, a process that potentially results in the production of N containing small molecules (Fig 9C; Table S2).

**Figure 9.**
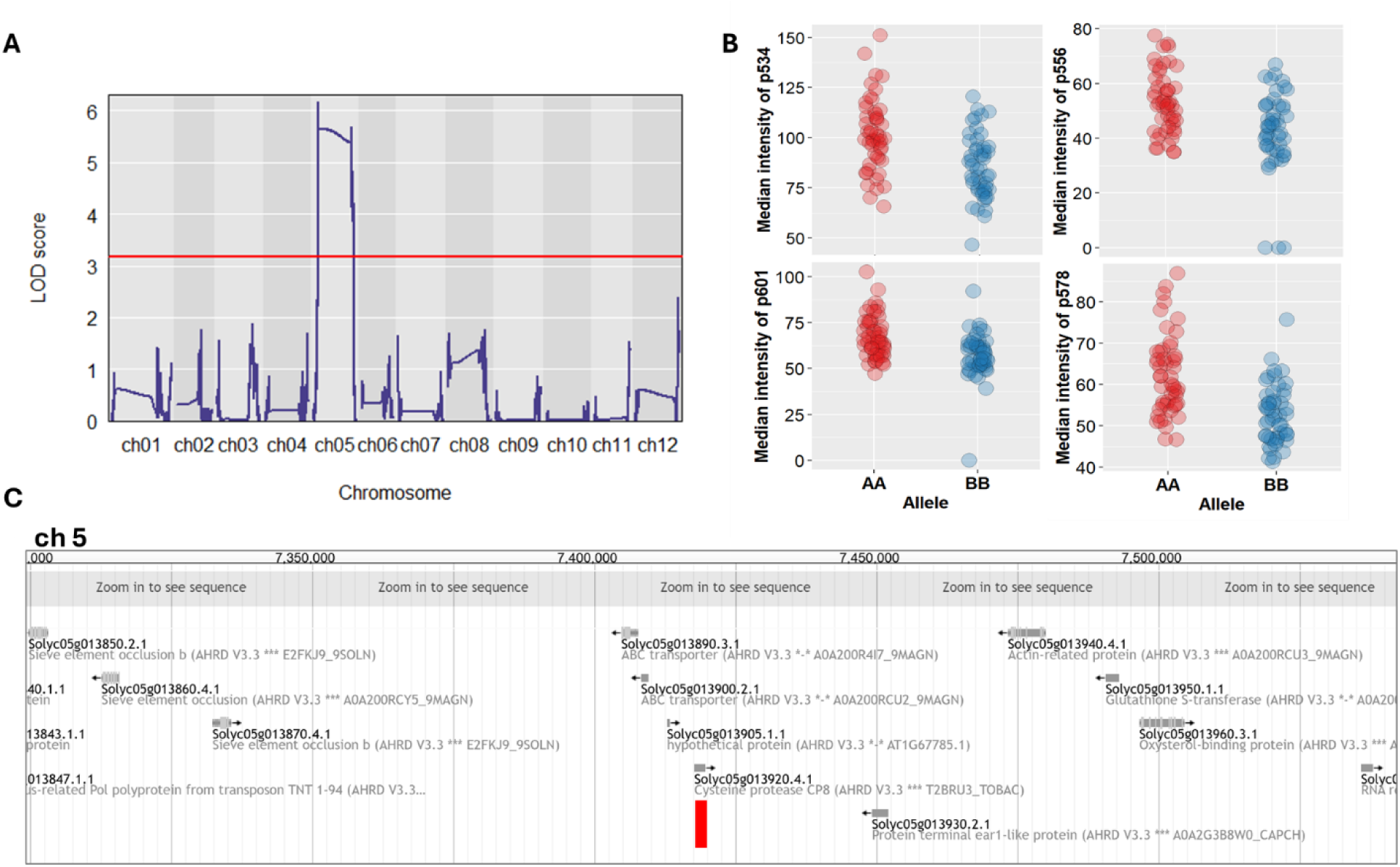
QTL associated with metabolic features p534, p556, p578, and p601. (A) Genome-wide scan showing the logarithm of odds (LOD) scores for the QTL. (B) Median intensity of p534, p556, p578, and p601 at the SNP marker located at 7.419 Mb on chromosome (ch) 5. Alleles are denoted as originating from *Solanum lycopersicum* cv. Moneymaker (AA) or *S. pimpinellifolium* (BB), illustrating parental allele contributions to metabolite variation. (C) Genomic region surrounding the SNP marker (highlighted with a blue line) visualized with JBrowse at solgenomics.net, based on gene model ITAG4.0.

### Overlap of metabolic and bacterial QTLs

Recently, Oyserman *et al*. (2022) reported significant QTLs associated with the abundance of specific bacteria and bacterial genes encoding proteins (i.e. metagenome assembly contigs) in the rhizosphere, using the same RIL population as used in the present study. To determine if there are metabolic features that potentially play a role in the recruitment of these microorganisms and/or microbial functions, the QTL list of the present study was interrogated for overlapping SNP markers with the study of Oyserman *et al*. This resulted in the identification of a single QTL which is associated with both a RE metabolic feature and the abundance of bacteria: p750 and ASV149 (Caulobacter) both associate with a SNP marker located at 81.990 Mb in Ch 1 having the same allele AA for both traits (Fig. 10; Table S3).

**Figure 10.**
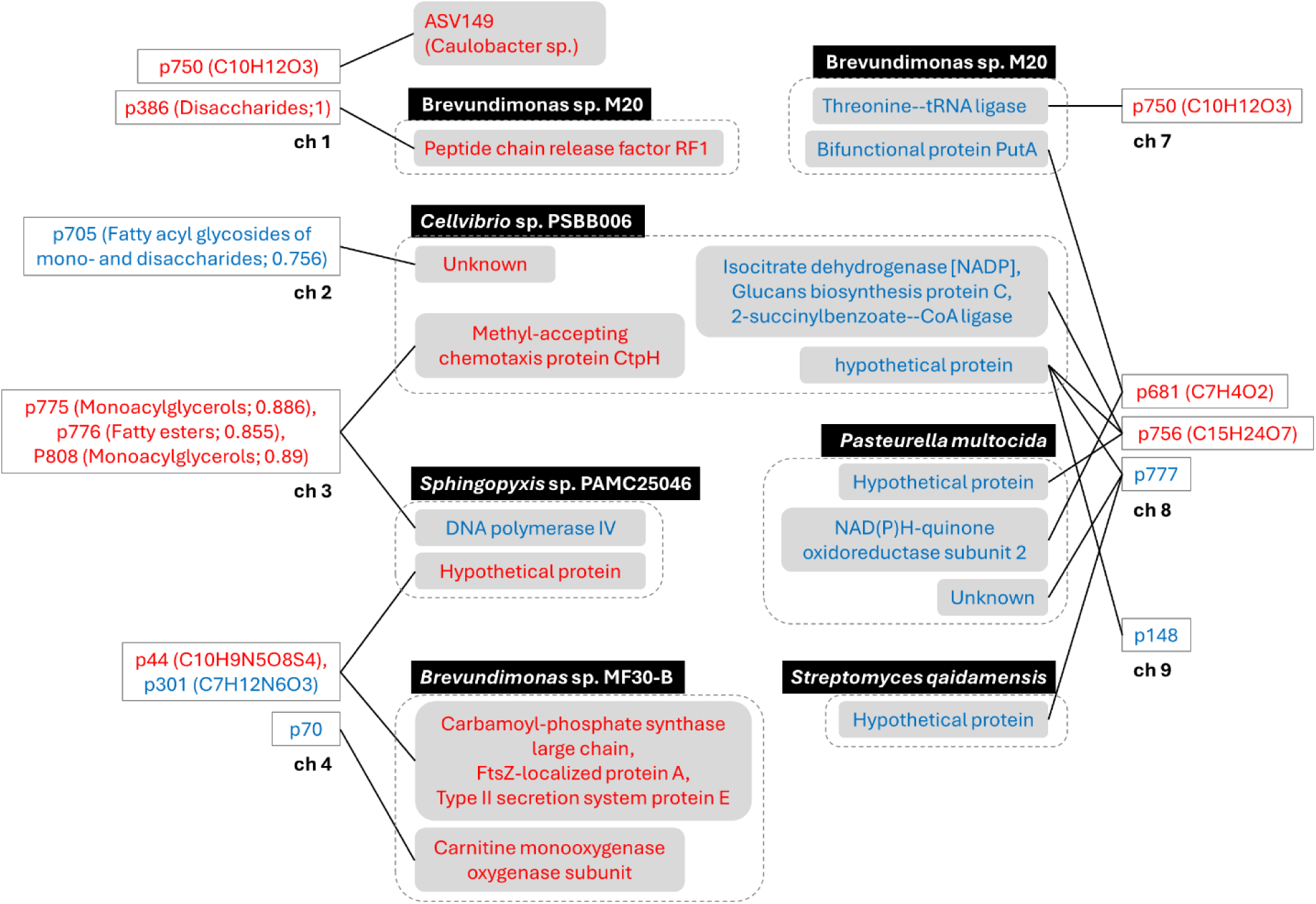
Overlapping QTLs between root exudate metabolic features (present study) and rhizosphere bacterial genome contigs (Oyserman et al., 2022) detected in the tomato RIL population. Traits shown in red are associated with alleles originating from *Solanum lycopersicum* cv. Moneymaker, while traits shown in blue are associated with alleles from *S. pimpinellifolium*.

In addition, 10 unique SNPs associated with 13 metabolic features overlapped with the QTLs for 15 bacterial genes (Table S3). Interestingly, this included 3 QTL-hotspots that linked to multiple metabolic features and bacterial genes (Fig. 10). For example, the SNP marker located at 2.053Mb in Ch 3 had an association with metabolic features p775, p776, and p808 (all predicted as fatty acids with high probability scores of 0.88, 0.85, and 0.89, respectively), and bacterial genes encoding ‘Methyl-accepting chemotaxis protein CtpH’ (originating from *Cellvibrio* sp. PSBB006) and ‘DNA polymerase IV’ (from *Sphingopyxis* sp. PAMC25046).

Another QTL-hotspot located at 62.615Mb on Ch 4 was shared by metabolic features p44 and p301 and bacterial genes encoding ‘Carbamoyl-phosphate synthase large chain’, ‘Type II secretion system protein E’ and ‘FtsZ-localized protein A’ (all from *Brevundimonas* sp. MF30-B) as well as a ‘Hypothetical protein’ (from *Sphingopyxis* sp. PAMC25046). The feature p44 is predicted to be derived from the alkaloid pathway with a score of 0.81, while p301 is linked to the fatty acid pathway with a probability score of 0.65. The QTL intervals for both metabolic features were relatively broad; however, several noteworthy candidate genes were identified in close proximity to the SNP marker (Fig. S1). These included a FAD-binding Berberine family protein (Solyc04g080395.1.1), an S-adenosyl-L-methionine-dependent methyltransferase superfamily protein (Solyc04g080360.4.1), and an NADPH-cytochrome P450 reductase (Solyc04g080340.3.1), all of which may contribute to the alkaloid biosynthetic pathway, as well as a 3-ketoacyl-CoA synthase (Solyc04g080450.1.1) with a potential role in fatty acid metabolism. Moreover, a gene encoding Pectinesterase (Solyc04g080540.3.1) near the SNP may influence bacterial colonization by modifying root cell wall pectin and affecting microbial attachment.

Lastly, the metabolic feature p756, associated with a SNP marker at 3.621 Mb on Ch 8, co-localized with several bacterial genes, including ‘2-Succinylbenzoate–CoA ligase’, ‘Isocitrate dehydrogenase [NADP]’, ‘Glucans biosynthesis protein C’, and a ‘Hypothetical protein’ (all from *Cellvibrio* sp. PSBB006), as well as a ‘Hypothetical protein’ from *Pasteurella multocida*. This overlap suggests that p756 influences on bacterial functions in the rhizosphere

Together, these results highlight that QTL mapping of tomato RE uncovers metabolic genetic associations. Importantly, several QTL hotspots were shared between RE metabolic features and bacterial genomic traits, found in the study by Oyserman (2022), suggesting that host genetic variation in exudate composition contributes to shaping the rhizosphere microbiome.

## Discussion

Despite the recognized importance of plant RE in plant fitness, many aspects of its composition and the genetic factors shaping it remain poorly understood. In this study, we provide the first genetic dissection of tomato RE using a metabolomics approach in a recombinant inbred line (RIL) population. We show that RE composition is strongly influenced by host genetics, with clear differences between wild and modern tomato species. Notably, we identified QTLs associated with the exudation of specific metabolites—including dihydroactinidiolide, 3-methylcrotonaldehyde, and chavicol—and QTLs that are linked to both metabolic features and bacterial functionality. To our knowledge, this is the first demonstration in tomato of a genetic link with RE composition that has been shaped by domestication, highlighting specific genomic regions that could be targeted in breeding to influence microbiome assembly and plant adaptation.

### The composition of tomato root exudate

In the past decade, research using metabolomics in biology has increased, as the metabolomics approach can convey large amounts of information that help us to understand complex natural phenomena between molecules, and this has been applied to the study of root exudates as well (van Dam and Bouwmeester 2016). Two earlier studies also report the composition of RE in tomato using metabolomics, to investigate the relationship between RE compounds and root-knot nematodes, and Fe and S deficiency (Yang et al. 2016; Astolfi et al. 2020). Several compounds that were found in the RE of these two studies were also detected in our study such as caffeine, benzaldehyde, and phosphoric acid, however, many of the compounds they report do not overlap with our RE profile (Table. S1). This is likely due to the low annotation rate of our RE profile (∼ 5%) and different methods for plant growth, root exudate collection, sample preparation, and chemical analysis method (Escolà Casas and Matamoros, 2021; McLaughlin *et al*., 2025). For instance, the two earlier studies collected RE by taking roots from growing substrate (e.g. soil) and immerging the roots in deionized or distilled water for a short period of time, while we collected RE by flushing the plants pots filled with soil. Advantages of our practice is that it more reflects the natural plant growing conditions and avoids root damage and osmotic stress associated with removing plants from soil and placing them in water (Williams *et al*., 2021). On the other hand, some exuded metabolites may have been lost through microbial degradation in the soil or adsorption to soil particles, as well as during the SPE concentration step due to selective retention by the column material. Conversely, we may also have gained metabolites originating from microbial production or from microbial conversion of plant-derived compounds. Indeed, we did not observe compounds commonly present in RE, such as coumarins, flavonoids, and strigolactones. Microbial production or conversion may also explain our low annotation rate because these metabolites are less likely present in most chemical libraries. Nevertheless, we could capture the differences in the RE composition between the two parents of the RIL population, and discover tomato genetic loci associated with a substantial number of metabolite features.

### p576 (dihydroactinidiolide)

The most intriguing metabolite feature in the RE, with a highly significant QTL, is p576, annotated as dihydroactinidiolide (dhA), with exceptionally high *H*^2^ and LOD score driven by its binary presence in RE of the two parents (Fig. 2, Fig. 4 and Table. 2). DhA is a β-carotene derived volatile terpene and created by oxidation of β-ionone (Havaux 2014). DhA seems to be involved in plant development; it inhibits germination in wheat, and increases lateral root branching in *Arabidopsis thaliana* (Kato et al. 2003; Dickinson et al. 2019). Moreover, it also plays an important role in photoacclimation. In *Arabidopsis* under high light exposure, the chemical oxidation of β-carotene by singlet oxygen (^1^O_2_) results in production of dhA. The accumulated dhA acts as a signaling molecule triggering ^1^O_2_ responsive gene expression that is necessary for plant adaptation to the high light intensity (Ramel et al. 2012; Shumbe, Bott and Havaux 2014). Several compounds belonging to apocarotenoids (e.g. β-cyclocitral, ABA, strigolactones) have been reported to have a function as signaling molecules (Moreno et al. 2021). Moreover, loliolide, the analogue of dhA, acts as a signaling chemical in the barnyard grass-rice allelopathic interaction in the rhizosphere (Li, Zhao and Kong 2020). All this suggests that dhA might be a potential rhizosphere signaling molecule.

Although dhA has been detected in the fruits of *S. lycopersicum* and *S. pimpinellifolium*, it has not previously been reported in their vegetative tissues nor in RE. In this study, we observed high levels of dhA in the RE, associated with the SNP marker originating from the parent SP (Fig. 4B). Notably, fruits of *S. pimpinellifolium* accumulate higher levels of dhA than cultivated *S. lycopersicum*, due to differences in the activity of 13 lipoxygenase TomLoxC (Gao et al. 2019). Whether TomLoxC or other apocarotenoid biosynthesis genes encoding carotenoid cleavage dioxygenase contribute to dhA variation in RE remains unclear; importantly, these genes do not co-localize with the QTL identified for dhA (p576) 0at 0.576 Mb of Ch 8 (Table 2; Fig. 4C). This QTL spans a very narrow interval harboring only 3 genes, including two terpene synthases, which partially belong to a larger terpene synthase (TPS) cluster comprising TPS18, 19, 20, 21, and 41 that produce mono- and di-terpenes, such as β-phellandrene and lycosantalene (Zhou and Pichersky 2020). The nucleotide sequences of this TPS cluster on Ch 8 are highly similar between *S. lycopersicum* and *S. pimpinellifolium*, except for few SNPs that do not affect functionality of the proteins (Matsuba et al. 2013). It is therefore unclear how this terpene synthase cluster is associated with dhA exudation.

One possible hypothesis is that the difference in dhA levels arises from variation in export activity rather than biosynthesis. Plant volatiles are transported across membranes by proteins such as ATP-binding cassette (ABC) transporters and non-specific lipid-transfer proteins (Adebesin et al. 2017; Liao et al. 2023). Although located just outside the QTL range for p576, an ABCG transporter gene (Solyc08g005580.3.1; Fig. 4C) lies very close to the QTL region and is highly expressed in root tissue (Ofori et al. 2018). In *Petunia hybrida*, the tissue-specific ABCG transporter PhABCG1 enables the passage of volatiles across the plasma membrane (Adebesin et al. 2017), raising the possibility that Solyc08g005580.3.1 contributes to dhA (p576) exudation in tomato roots. The CDS of this gene is identical between *S. lycopersicum* and *S. pimpinellifolium* LA1589 (BLAST search at solgenomics.net; data not shown), suggesting that any functional difference, if present, may stem from transcriptional regulation rather than protein sequence variation.

### P747 (3-methylcrotonaldehyde)

RE metabolite feature p747 had 2 significant QTLs (at 81.946 Mb on Ch 1 and at 3.092 Mb on Ch 7; Fig. 5) and is also annotated as a volatile, 3-methylcrotonaldehyde (also known as 3-methyl-2-butenal). 3-Methylcrotonaldehyde has been reported in tomato fruits (Encinas-Basurto et al. 2017; da Silva Souza et al. 2020), and it is synthesized through oxidation of isopentenyl adenosine and N6-dimethylallyladenine by cytokinin (CK) oxidases, resulting in CK deactivation (Mok and Mok 2003). The role of 3-methylcrotonaldehyde in the rhizosphere is unknown, but the precursor CK synthesized in roots seems to be engaged in events in the rhizosphere. For instance, zeatin in tomato root exudate attracts root-knot nematode *Meloidogyne incognita* (Kirwa et al. 2018). Both QTL regions, do not contain CK oxidases or CK responsive genes, of which the expression may increase or decrease upon induction of the CK pathway (Gupta et al. 2013). However, the region around the p474-associated QTL marker at Ch 1 did contain genes encoding proteins involved in plant stress response regulation that might link to the CK pathway (Fig 5C): Acyl-CoA-binding domain-containing protein 4 (Solyc01g099350.3.1), calcium-dependent lipid-binding family protein (Solyc01g099370.3.1), and WD-40 repeat containing protein (Solyc01g099400.4.1).

The Acyl-CoA-binding domain-containing protein (ACBP) family is central to lipid metabolism, participating in lipid binding, transport, and vesicle trafficking across many organisms (Qiu and Zeng 2020). Although their roles in tomato remain poorly understood, studies in *Arabidopsis* suggest that ACBPs interact closely with hormone signaling pathways, including cytokinin (CK), and help plants adapt to environmental stress by altering membrane lipid composition (Du et al. 2016) (Abdelrahman *et al*., 2021). Given this, the ACBP4 gene near the p474-associated QTL marker on Ch 1 may play a role in CK-linked lipid remodeling.

Calcium-dependent lipid-binding protein (CLB) family proteins sense elevated Ca^2+^ signals upon abiotic stresses in plants and play a role in regulating the response of plants to the stresses including drought and salinity (De Silva et al. 2011; Gao et al. 2020). Interestingly, the region around the other p747-associated QTL at 3.092 Mb on Ch 7 harbors a gene encoding a calcium-transporting ATPase (Solyc07g008320.3.1), possibly linked to the CLB (Solyc01g099370.3.1) that we found on Ch 1.

WD-40 repeat (WDR) proteins, known for their β-propeller structure, are versatile regulators of protein interactions and transcriptional control in plants (Miller, Chezem and Clay 2016). They participate in diverse processes such as cell wall development, flavonoid biosynthesis, and defense signaling (Guerriero, Hausman and Ezcurra 2015; Miller, Chezem and Clay 2016; Gao et al. 2018). Intriguingly, the QTL region on Ch 7 also harbors a gene encoding 4-coumarate:CoA ligase (Solyc07g008360.2.1), which converts 4-coumarate into 4-coumaroyl-CoA, a central intermediate in the phenylpropanoid pathway. Since CK has been shown to regulate flavonoid biosynthesis (Abdelrahman *et al*., 2021), the proximity of WDR proteins and 4-coumarate:CoA ligase raises the possibility of an integrated CK–flavonoid regulatory link within this QTL.

We blasted the coding/protein sequence of genes discussed above in *S. pimpinelifolium* LA1589 using BLAST (solgenomics.net), and the nucleotide sequences were identical with *S. lycopersicum*, and even if not, the protein sequences were the same (data not shown). Therefore, it is possible that the quantitative differences of p747 in tomato RE was due to unknown transcriptional regulation. Moreover, due to the wide range of p747-associated QTL intervals (80.722 – 86.859 Mb for Ch 1 and 2.271 – 57.870 Mb for Ch 7), only the genes located at 0.5 Mb before and after the SNP marker are discussed here; we cannot exclude that other genes located further away, within the interval, are underlying the variation in p747 concentration in the RE.

### P613 (Chavicol)

Metabolic feature p613 in the RE had a significant QTL at 60.626 Mb in Ch 7 (Table 2; Fig. 6). This feature was annotated as chavicol, a phenylpropanoid. Chavicol is synthesized by chavicol/eugenol synthase (CVS/EGS) from p-coumaroyl acetate, and CVS/EGSs in plants have been discovered in basil, strawberry, petunia, *Clarkia breweri*, creosote bush and anis (Atkinson 2018), but not yet in tomato. The methylated form of chavicol, methyl chavicol (also known as estragole), has been demonstrated to play a role in the interaction of plants with other organisms. For example, methyl chavicol displayed antimicrobial activity to several bacteria and fungi in vitro (Lachowicz et al. 1998; Suppakul et al. 2003), and insecticidal activity to corn pest *Spodoptera frugiperda* (de Menezes et al. 2020). Considering its biological activity it might also influence the soil microbial biodiversity. Methyl chavicol is formed from chavicol by an O-methyltransferase that uses S-adenosylmethionine as the methyl donor (Yauk et al. 2015; Atkinson 2018). A chavicol O-methyltransferase has not yet been discovered in tomato. Interestingly, however, Solyc07g052110.4.1 is located close to the p613-associated SNP marker at 60.568 Mb in Ch 7 and encodes a S-adenosyl-L-methionine-dependent methyltransferases superfamily protein (Fig. 6C). Further evaluation is needed to confirm if this protein catalyzes methyl chavicol production, but if this is the case, this protein might contribute to the different amount of p673 in the RE across the RIL population. This is not due to sequence polymorphism, as BLASTing of the *S. lycopersicum* Solyc07g052110.4.1 nucleotide sequence against the *S. pimpinelifolium* LA1589 genome showed they are identical (solgenomics.net; data not shown). Possibly, there is a polymorphism in a cis-regulatory element resulting in a difference in transcription, within the QTL region.

### Terpenoids

There were many other metabolite features significantly associated with QTLs, though their exact identities remain unknown. SIRIUS prediction did predict two metabolic features to belong to the terpenoids, and both of them are linked to intriguing QTLs.

Metabolic feature, p666, had a high probability score (0.734) of being a C30 triterpenoid and was significantly associated with a QTL marker at 67 Mb on Ch 9 (Table 2; Fig. 7). This marker is adjacent to a gene encoding an ABC transporter Pleiotropic Drug Resistance (PDR) protein (Solyc09g091660.3.1; Fig 7C). It is tempting to speculate that this transporter may be involved in the transport of p666, and that a genetic polymorphism results in differences in p666 intensity in the RE between the two tomato parents. There is evidence supporting the role of PDR transporters in terpenoid transport. For instance, PgPDR3 in *Panax ginseng* contributes to C30 triterpenoid ginsenoside accumulation under methyl jasmonate induction (Zhang et al. 2013), while NpPDR1 (formerly NpABC1) in *Nicotiana plumbaginifolia* facilitates the secretion of the antifungal diterpene sclareol in response to microbial pathogens (Jasiński et al. 2001; Stukkens et al. 2005). Similar evidence has been shown with AtPDR12 in *Arabidopsis thaliana*, emphasizing the role of PDR transporters in sclareol transport and plant defense (Campbell et al. 2003). Interestingly, Solyc09g091660.3.1 shares high similarity with AtPDR12 (99% query cover, 71.82% identity in NCBI protein BLAST; data not shown) and is highly expressed in root tissue (based on Evorepro database). Moreover, it is upregulated during effector-triggered immunity at both transcript and protein levels (Bashary et al. 2024). This supports its potential role in terpenoid secretion and defense responses. There was no polymorphism in the CDS of Solyc09g091660.3.1 or its protein sequence between the two parents BLAST search at solgenomics.net; data not shown). Therefore, if Solyc09g091660.3.1 is indeed involved in p666 exudation, its effect is likely attributable to differences in transcriptional regulation.

Another metabolite feature, p634, was also predicted to be a triterpenoid (probability 0.55) and was significantly associated with a QTL marker at 63.245 Mb on Ch 4 (Table 2; Fig. 8). This marker is located adjacent to genes involved in auxin signaling (Fig. 8C), including Auxin Response Factor 5 (Solyc04g081240.2.1), Small Auxin Up-Regulated RNA51 (Solyc04g081250.1.1), and Small Auxin Up-Regulated RNA52 (Solyc04g081270.1.1). Evidence supporting a role for auxin in triterpenoid biosynthesis is sparse. Although IAA reduces solanoeclepin A accumulation in tomato hairy roots in vitro (Shimizu *et al*., 2023), any direct involvement in the p634 pathway remains speculative.

### Nitrogen containing metabolites

Nitrogen containing metabolites such as amino acids and peptides have been found in the RE of a wide range of plants, and suggested to have diverse ecological functions in the rhizosphere (Tan et al. 2024). We also detected metabolic features predicted to contain nitrogen in the RE of the RIL population and associated QTLs. From the latter, an interesting QTL hotspot is located at 7.418 Mb on Ch 5 (Table 2; Fig9), showing association with 4 metabolite features predicted to contain nitrogen (i.e. p534, p556, p578, and p601). The SNP associated with these metabolic features is located next to a gene encoding Cysteine Protease 8 (Solyc05g013920.4.1; Fig. 9C), an enzyme that cleaves peptide bonds in proteins. This enzymatic activity might influence the composition of nitrogen-containing molecules in root exudates, potentially affecting chemical communication in the rhizosphere.

### RE-Microbiome related QTLs

In the present study, we demonstrated that the composition of the tomato RE is shaped by genetic factors, resulting in substantial differences in the RE between wild and cultivated tomato (Fig. 1). These differences potentially drive the differences in the root-associated microbiome among the genotypes within this RIL population as demonstrated before (Oyserman et al. 2022). Indeed, we detected overlapping QTLs between the abundance of metabolic features in the tomato RE and bacterial ASVs and functionality in the rhizosphere as identified by Oyserman *et al* (Table S3).

An intriguing overlapping QTL at 62.61 Mb on Ch 4 is associated with metabolic features p44 and p301 and several *Brevundimonas* genes, including those encoding ‘Carbamoyl-phosphate (CP) synthase large chain’, ‘FtsZ-localized protein A’, and ‘Type II secretion system protein E’ (Fig. 10; Table S3). This QTL region is particularly notable due to the presence of multiple genes that may influence plant metabolism and, hence, microbial abundance in the rhizosphere. Metabolic feature p301 was predicted to be derived from the fatty acid pathway with a probability score of 0.65, while p44 was linked to the alkaloid pathway with a score of 0.81. Several nearby genes are known to facilitate these pathways (Fig. S1). For alkaloid metabolism, FAD-binding Berberine family protein (Solyc04g080395.1.1), S-adenosyl-L-methionine-dependent methyltransferase superfamily protein (Solyc04g080360.4.1), and NADPH-cytochrome P450 reductase (Solyc04g080340.3.1) may contribute, given their established roles in various alkaloid biosynthetic pathways (Daniel et al. 2017; Chakraborty et al. 2023; Lee et al. 2024). Solyc04g080450.1.1, encodes a 3-ketoacyl-CoA synthase, which is involved in fatty acid metabolism. This enzyme primarily elongates long-chain fatty acids (C16–C34; Huang et al. 2023), making its involvement in p301 biosynthesis—predicted to be a C7 or C9 fatty acid derivative—not entirely intuitive. Instead, it may influence p301 levels indirectly by regulating long-chain fatty acid degradation, which could impact precursor availability or metabolic flux.

Beyond its potential role in metabolite biosynthesis, this QTL region may also influence microbial interactions. The presence of a QTL for *Brevundimonas*-associated genes suggests that these metabolites contribute to the recruitment or growth of this bacterium. Bacterial ‘CP synthase’, for example, catalyzes the synthesis of CP, a precursor for pyrimidine and arginine—key components of nucleic acids and proteins (Thoden et al. 2002). Similarly, ‘FtsZ-localized protein A’ is essential for bacterial cell division (Lariviere et al. 2018). Finally, while metabolic features in the RE likely play a crucial role in shaping the microbiome, additional genes in the QTL region – not related to metabolism - may also play a role in mediating the plant-microbe interaction.

One such gene in the QTL region is Solyc04g080530.3.1, encoding ‘Pectinesterase’, an enzyme that modifies pectin in the plant cell wall. Pectinesterase activity enhances plant defense by generating oligogalacturonides, which act as damage-associated molecular patterns, and by regulating cell wall integrity, limit microbial invasion (Lionetti et al. 2017). Notably, some microbes—including both pathogens and beneficial symbionts—actively modulate pectinesterase activity to facilitate their colonization (Su, 2023). This highlights the potential role of Solyc04g080530.3.1 in shaping root-associated microbial communities in response to environmental and metabolic cues.

Another notable overlapping QTL linking metabolic features and microbial function is located at 2.053 Mb on Ch 3 (Fig. 10; Table S3). This QTL is associated with p775, p776, and p808, as well as a bacterial gene encoding the “Methyl-accepting chemotaxis protein CtpH” from *Cellvibrio* sp. PSBB006. In *Pseudomonas* spp., CtpH senses fluctuations in inorganic phosphate (Pi) and guides chemotaxis toward higher Pi concentrations (Wu et al. 2000; Rico-Jiménez et al. 2016)(Wu et al. 2000; Rico-Jiménez et al. 2016), suggesting that tomato genotypes carrying this QTL may influence Pi dynamics in the rhizosphere via recruitment of bacteria carrying this gene. The associated metabolite features, p775, p776, and p808, were annotated as fatty acids with high probability scores. While their direct role in Pi availability is unclear, fatty acids are key components of bacterial cell walls, and *Cellvibrio* sp. PSBB006 may utilize these compounds for growth or colonization.

## Conclusion

Our findings add to a growing body of work demonstrating that root exudation is not only a biochemical process but also a genetically regulated trait with downstream ecological consequences. Our work extends these insights to tomato, revealing specific QTLs that govern the root exudation of potentially ecologically relevant compounds. This has important implications for agriculture and breeding: tailoring crop varieties to exude beneficial metabolites could provide a sustainable means of enhancing stress tolerance, nutrient acquisition, and disease resistance by steering microbiome assembly. Future research should focus on functional validation of the candidate genes and metabolites identified here, as well as their direct effects on microbial recruitment and plant performance, under controlled and, subsequently, field conditions. Incorporating high-resolution metabolomics, microbial profiling, and gene-editing approaches will be critical for establishing causal links. Overall, our results highlight the potential of integrating metabolomic and genetic knowledge into breeding programs to not only improve crop productivity but also contribute to environmentally sustainable agriculture by harnessing natural plant–microbe interactions.

## Acknowledgements

We would like to thank Alessandra Guerrieri for helping the greenhouse experiment and Marc Galland for helpful suggestions. We acknowledge funding from the European Research Council (ERC) Advanced grant CHEMCOMRHIZO 670211, the Dutch Research Council (NWO/OCW) Gravitation programme Harnessing the second genome of plants (MiCRop) and the University of Amsterdam Research Priority Area Systems Biology (SysBA).

---

**Figure S1.**
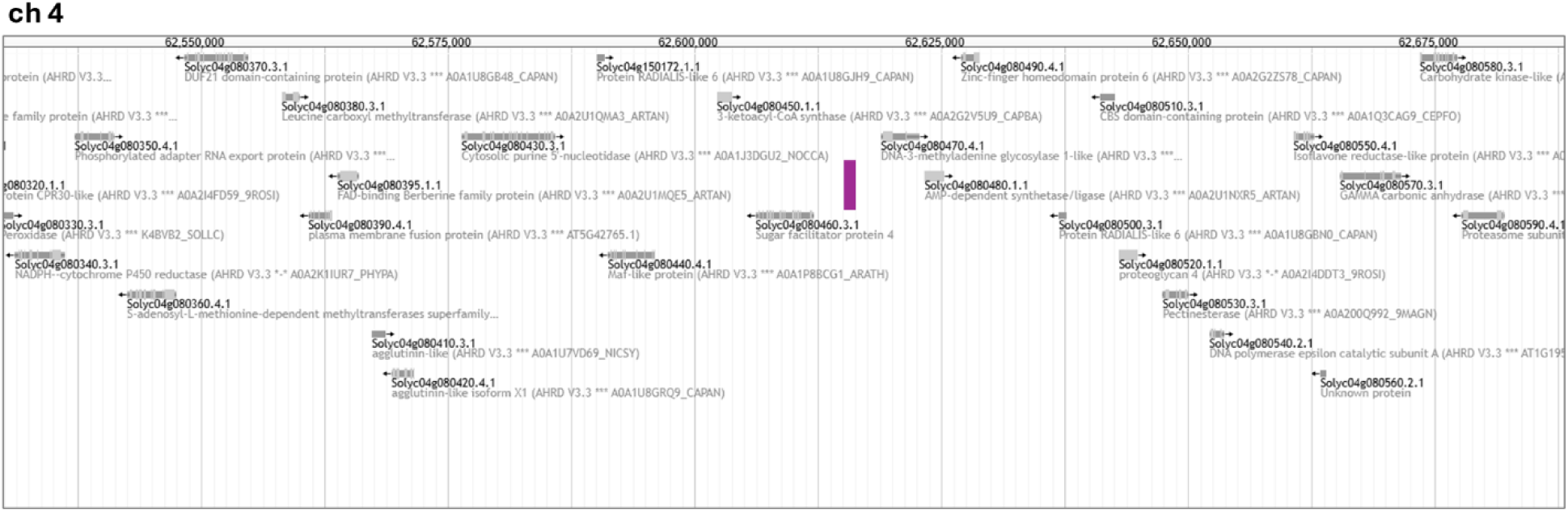
Genomic region surrounding the SNP marker (located at 62.615 Mb on chromosome 4; highlighted with a purple line) associated with the metabolic features p44 and p301. Several microbial genes were also mapped to this marker, including *carbamoyl-phosphate synthase large chain*, *type II secretion system protein E*, and *FtsZ-localized protein A* (all from *Brevundimonas* sp. MF30-B), as well as a *hypothetical protein* (from *Sphingopyxis* sp. PAMC25046). The region was visualized in JBrowse at solgenomics.net using the ITAG4.0 gene model.

